# AI Analysis of a Copy Number Variant Database Identifies a Genetic Factor for a Murine Model of the Metabolic Syndrome

**DOI:** 10.64898/2026.08.05.743102

**Authors:** Wenlong Ren, ZhuanFen Cheng, Gary Peltz

## Abstract

Copy number variants (CNVs) are a major source of genetic diversity and could contain some of the missing heritability for mouse models of human disease. However, mouse CNVs have not been comprehensively characterized because they are difficult to resolve in repeat-rich, segmentally duplicated or reference sequence-absent regions of the genome. Here we analyzed long range sequence (LRS) data for 40 inbred mouse strains and characterized CNVs using pangenome graph-based (and other) methods and a C57BL/6J telomere to telomere (T2T) genome reference sequence. We resolved 1,594 high-confidence CNVs that often overlap tandem repeats (60.3%), segmental duplications (44.8%) or pericentromeric regions (11.5%); and 131 CNVs were T2T sequence-specific. CNVs affected 384 protein-coding genes, which spanned a range of important functional classes. The 40-strain pangenome map expanded the genome sequence from 2.29 to 3.32 Gb, with the wild-derived strains accounting for the largest sequence increments. Two different AIs were sequentially used to analyze this database and identify a 29-kb deletion CNV within the *Nlrp1b* locus of KK mice that contributed to the metabolic syndrome they develop. Human *NLRP1* alleles also were associated with metabolic syndrome features in human populations. Hence, AI analyses of this comprehensive T2T pangenome-based resource could uncover some of the missing heritability for mouse models of human diseases and biomedical traits.

---

The laboratory mouse has been the premier model organism for biomedical research. Mouse models have been used to discover or test many of today’s commonly used therapies ^1,2^ and to characterize genetic factors for human diseases ^3–5^. Our ability to identify causative murine genetic factors depends upon having a comprehensive map of genetic variation among the inbred strains. While SNPs and indels have been extensively characterized ^6–8^, the missing heritability for several mouse models were identified by analysis of other types of genetic variants. For example, genetic factors for mouse models, which had not been found over a preceding 50-year period, were identified by analyzing recently generated structural variant (SVs) ^9^ and tandem repeat (TR) ^10^ databases. Some of the missing heritability for the 3500 phenotypes measured in inbred strain panels, many of which reflect human disease traits ^11,12^, could be identified by analysis of copy number variants (CNVs) (i.e., changes in the number of copies of a genomic DNA segment), which are a major source of genetic diversity. Non-allelic homologous recombination between repeated sequences produces recurrent CNVs with the same breakpoints in different individuals, while CNVs that arise from non-homologous recombination mechanisms have variable endpoints ^13^. Due to their size (ranging from kilobases to megabases), recurrent CNVs account for a larger fraction of the total base pair differences between genomes than SNPs ^14,15^. CNVs have a profound influence on chromatin structure, gene regulation, development, and disease susceptibility ^13^. Mouse and human CNVs contribute to congenital anomalies, multiple complex human traits ^16–18^, hearing loss ^19,20^ and neurodevelopmental conditions ^21^.

Earlier efforts identified clusters of CNVs in the mouse genome ^22–25^, but the diversity and distribution of large CNVs could not be fully defined because they often occur within complex genomic regions, which are difficult to resolve using linear assemblies ^26–28^ or array-based hybridization methods. Therefore, we combined three state-of-the-art variant discovery strategies to improve our ability to analyze CNVs in these regions. (i) The use of long read sequencing (LRS) that produces reads of length ≥20 kb improved our ability to analyze complex genomic regions ^29–31^, which enables CNV breakpoints to be precisely determined ^32^. (ii) Pangenome analysis overcomes the inherent limitations of using a single reference sequence. A pangenome graph for a species displays the DNA sequences shared by all individuals along with those that are unique to one or more individuals. Multiple assemblies are integrated into a unified graph that represents alternative haplotypes, non-reference sequence, presence/absence variation (PAV) and multi-copy structures ^33–35^. A linear reference sequence often collapses divergent or duplicated regions, while pangenome graphs preserve population-level structural diversity and more faithfully represents the complex genomic configurations present in individuals. Pangenome graphs have revealed previously hidden sequence variation for humans and plants; have markedly expanded human CNV (and other large SV) catalogues ^36–39^; and have been used to characterize haplotype divergence and TR diversity across mouse subspecies ^40^. They provide a powerful framework for resolving non-reference sequences and for dissecting structural diversity with high resolution. (iii) Human telomere-to-telomere (T2T) sequencing has resolved previously inaccessible genomic regions and provides a solid foundation for high-resolution CNV and SV discovery ^41,42^. T2T assemblies for plants, ruminants and apes have identified CNVs that were not resolved using legacy reference sequence ^43–46^. Mouse genome T2T reconstructions revealed strain-specific differences in centromeric satellites, telomeric architectures and duplicated gene clusters that were missed using standard CNV pipelines ^47,48^. Since T2T sequences has had a transformative effect on large variant discovery, we use a pangenome graph and high-fidelity LRS to establish a T2T-anchored framework for large-scale CNV discovery across 40 inbred mouse strains.

We demonstrate the utility of this CNV atlas by analyzing a widely studied polygenic model for diet-induced obesity, insulin resistance and type 2 diabetes (T2D) (i.e., the metabolic syndrome) that develops in KK mice ^49–51^. A few mutations in KK and its sub-strains contribute to components of its metabolic syndrome. The Lethal Yellow (A^y^) mutation on the KK background (KK-A^y^ mice) induces hyperphagia and an early onset and severe metabolic syndrome ^52^ and a KK Apolipoprotein A-II variant (*Apoa2^b^* allele) elevates total- and HDL-cholesterol ^53,54^. Although KK mice have been studied for over 64 years ^49^, the major genetic drivers of its metabolic syndrome have not been identified. We demonstrate the utility of this database for genetic discovery by sequentially using two AIs (‘*stacked AI method’*) to analyze the CNV database and identify a CNV that contributes to the KK metabolic syndrome. Moreover, we demonstrate that mouse genetic results can guide the search for causative human genetic factors by examining associations between alleles in a homologous human gene with a human disease that resembles the mouse pathology.

## Results

*Overview*. The complete C57BL/6J T2T genome ^48^ was used as the backbone for this analysis because it has 213 Mb of additional sequence, 225 new genes and 87 filled gaps relative to the C57BL/6J GRCm39 reference sequence. Genomic LRS sequencing data was generated using a PacBio Revio instrument with the HiFi system for 36 classical and four wild-derived inbred strains, which had an average of 88.5 GB (30x coverage) per strain ^9^. The full names and abbreviations used for each strain analyzed, and the sequencing parameters are indicated in **Fig. S1** and **Table S1**. Three complementary strategies were optimized for detecting CNVs (≥10 kb): (i) graph-based CNV discovery used the Minigraph-Cactus pangenome pipeline ^55^, which quantified copy number states across homology blocks; (ii) assembly-based CNV detection by aligning the HiFiasm ^56,57^ sequence assemblies to the T2T reference, and variants were identified using SVIM-asm ^58^ and PAV ^59^; and (iii) Sawfish ^60^ was used to identify CNVs based upon read depth and structural breakpoints (**Fig. 1**).

**Figure 1.**
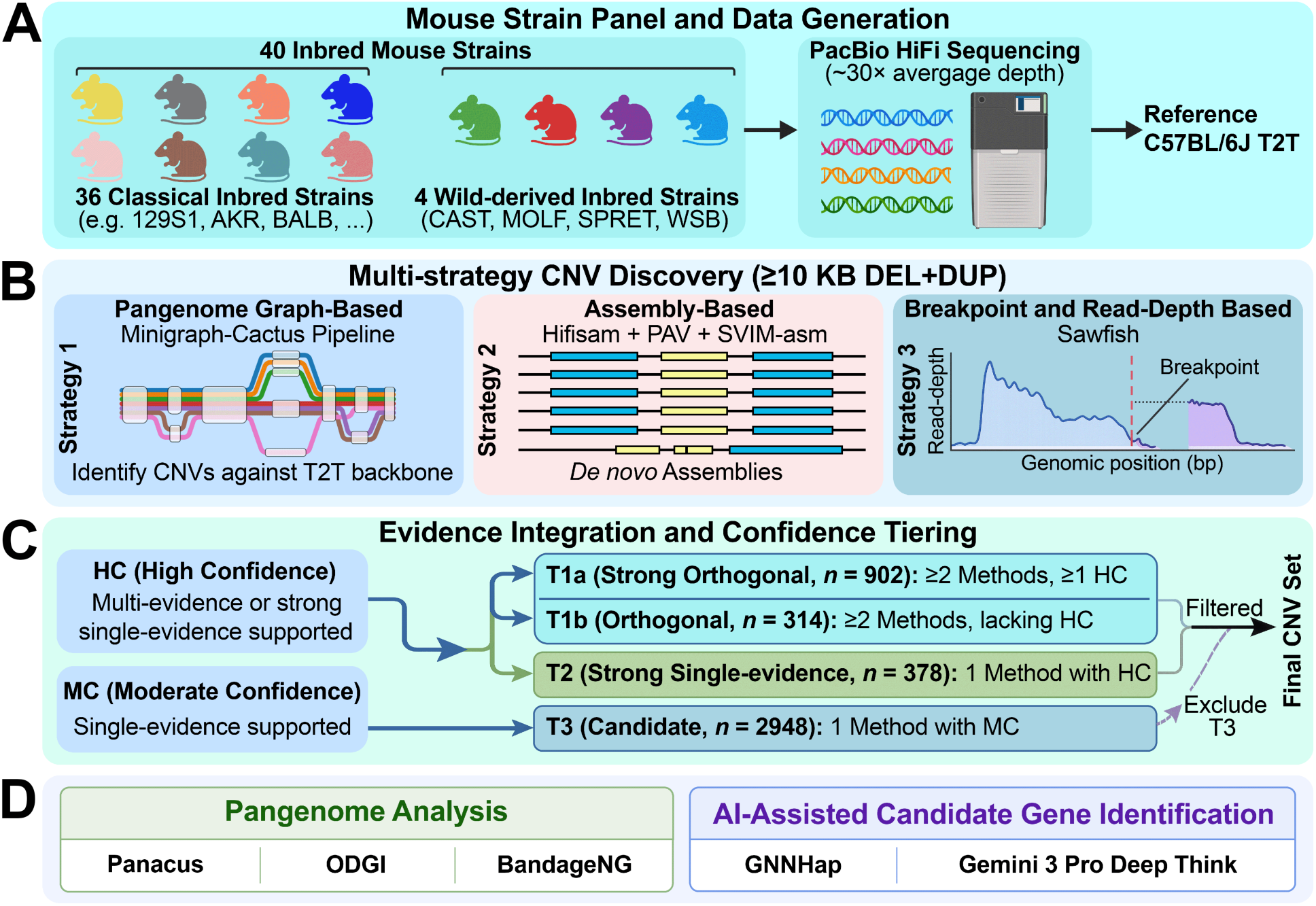
The methods used to characterize CNVs in the genomes of 40 inbred strains. (**A**) LRS was performed on 40 inbred mouse strains and the C57BL/6J T2T assembly was used as the reference. (**B**) Genomic LRS data from 40 strains was analyzed using 3 complementary CNV detection strategies: (i) graph-based CNV discovery used the Minigraph-Cactus pangenome pipeline, which quantified copy number states across homology blocks; (ii) assembly-based CNV detection by aligning HiFiasm sequence assemblies to the T2T reference, and variants were identified using SVIM-asm and PAV; and (iii) Sawfish was used to identify CNV based upon read depth and structural breakpoints. (**C**) The confidence level for each reported CNV was assessed based on both the number of supporting methods and the confidence categories assigned by each method. CNVs were then stratified into the tiers shown here according to these criteria. (**D**) Whole-genome pangenome analyses were performed using Panacus, ODGI, and BandageNG to characterize pangenome architecture and sequence diversity across the 40 mouse strains. Candidate genes associated with a phenotype of interest were identified using an AI-assisted framework that included GNNHap and Gemini 3 Pro DeepThink.

To assemble the database, we used a ≥10 kb CNV size threshold for three reasons. (i) It distinguishes this analysis from our prior characterization of structural variants (≥ 50 bp) ^9^, removes mobile element insertions (e.g., L1 retrotransposons) and focuses on large-scale genomic rearrangements. (ii) It aligns with the resolution boundaries used by Chromosomal Microarray Analysis, which enables a direct comparison with established cytogenetic benchmarks. (iii) Most significantly, large CNVs are more amenable to direct sequence-resolved characterization, which reduces the reliance on indirect read-depth approximations that are often unreliable in repetitive or structurally complex genomic regions. Instead, we utilize pangenome, assembly-based and alignment-based tools (Minigraph-Cactus, PAV, SVIM-asm, and Sawfish) to characterize large genomic alterations—including those within complex segmental duplications— at base pair resolution.

### CNV database

The CNVs were classified into tiers based upon the discovery methods used for their identification and the confidence levels provided by the supporting evidence. CNVs identified by at least 2 different methods were labelled as Tier1, and those identified by one method with high confidence were Tier 2. While candidate CNVs (Tier 3), which were identified by only one method with moderate confidence, were not further considered here. By these criteria, 1594 CNVs were identified with high confidence (Tiers 1a, 1b and 2) in the 40 strains with base pair resolution (**Fig. S2** and **Supplemental Data File S1**). Tier1a,1b, and 2 contained 902 (901 deletion + 1 duplication), 314 (306 deletion + 8 duplication), and 378 (370 deletion + 8 duplications) CNVs, respectively. The four wild-derived strains exhibited substantially higher CNV counts than the classical laboratory strains; SPRET had the highest number of CNVs (733 CNVs), followed by CAST (501), MOLF (475), and WSB (248) (**Fig. 2A**). In contrast, several of the closely related classical strains (B6J, B10J, and B10D2) had very few CNVs. Most CNV alleles were shared by only one (823), two (207) or three (153) CNVs, respectively, and the number of shared CNVs progressively decreased as the number of strains sharing that CNV increased (**Fig. 2B**). The CNVs ranged in size from 10,006 bp to 341,465 bp. CNVs in the 4 wild-derived strains were significantly shorter than those in the classical inbred strains, with median lengths of 13.5 kb and 14.9 kb, respectively (two-sided Wilcoxon rank-sum test, *P* = 2 x 10^-22^) (**Fig. 2C**). Many CNVs were associated with complex genomic regions within 5 kb of their breakpoints, which included those with tandem repeats (TR, 60.3%), segmental duplications (SD, 44.8%) and pericentromeric regions (Peri, 11.5%) (**Fig. 2D**). Phylogenetic clustering and a pairwise similarity heatmap based upon CNV alleles present in 40 inbred mouse strains are consistent with the known strain phylogeny ^61,62^: the four wild-derived strains (MOLF, CAST, SPRET, WSB), DBA1 and DBA2, and the NZW, NZB, and NZO strains formed separate clusters (**Fig 2E**).

**Figure 2.**
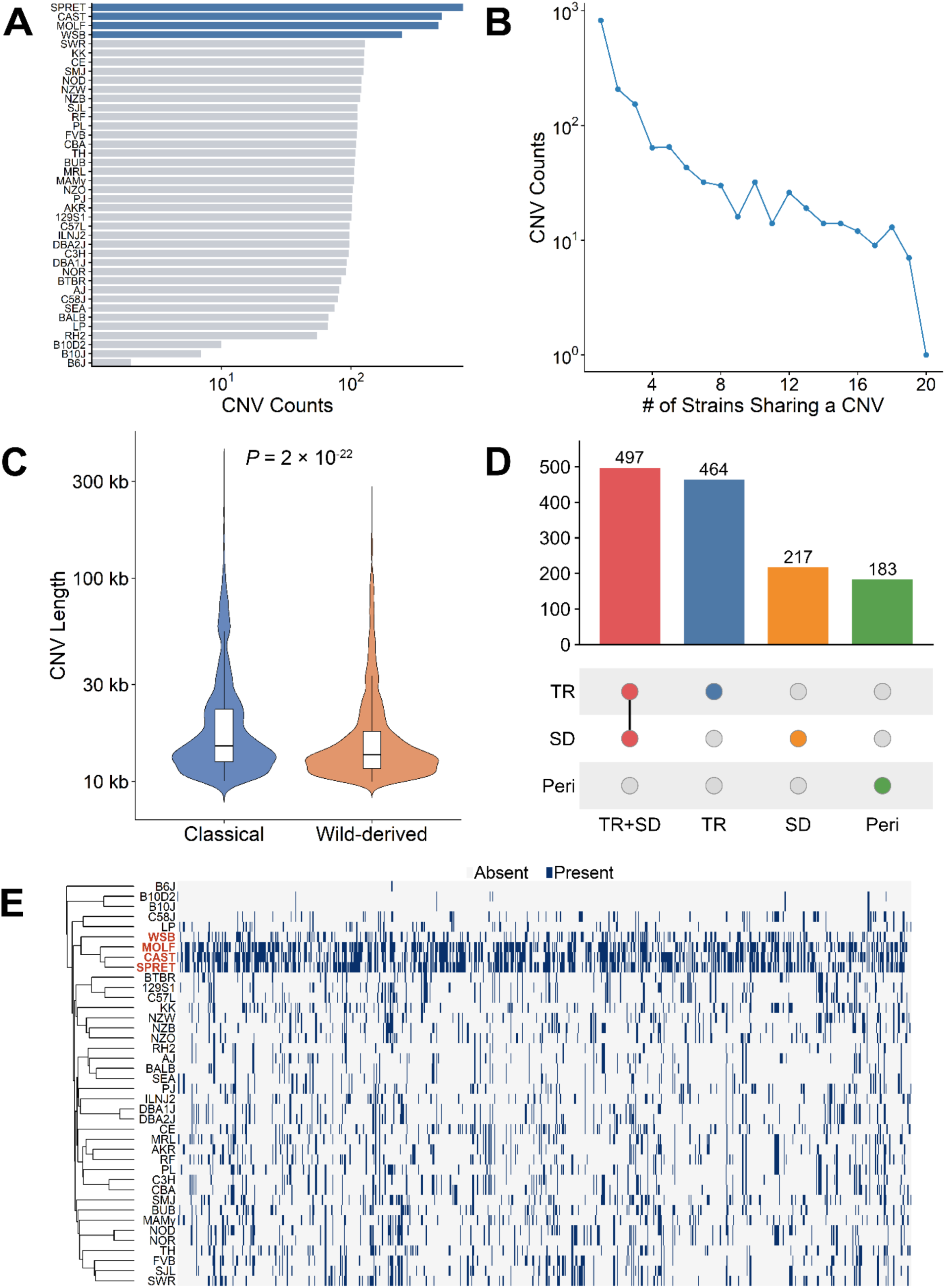
The distribution and allele sharing pattern of CNVs across 40 mouse inbred strains. (**A**) The total number of CNVs identified in each of the 40 mouse inbred strains. The CNV counts are shown on a log10 scale to facilitate comparison across strains with highly variable numbers of CNVs. The four wild-derived strains (SPRET, CAST, MOLF, WSB) had the largest number of CNVs relative to the classical inbred strains, which is consistent with their increased genetic divergence from the C57BL/6J reference genome. (**B**) The distribution of CNV allele sharing across the 40 mouse inbred strains. The x-axis indicates the number of strains sharing a given CNV, whereas the y-axis represents the number of CNVs on a log10 scale. Most CNVs were rare and observed in only one to three strains, indicating that the majority of CNVs in the mouse population are strain-specific or low-frequency. (**C**) The size distributions of CNVs in the 36 classical and four wild-derived mouse strains (MOLF, CAST, SPRET, and WSB). The y-axis represents CNV length on a log10 scale. Violin plots show the density distribution of CNV lengths, with wider regions indicating higher CNV frequency. The embedded boxplots indicate the interquartile range (IQR; 25th–75th percentiles), and the horizontal line within each box denotes the median CNV length (Classical: 14.9 kb; Wild-derived: 13.5 kb). Whiskers extend to 1.5 × IQR. The statistical significance of the difference in their size distributions was assessed using a two-sided Wilcoxon rank-sum test (*P* = 2 × 10^-22^). (**D**) An UpSet plot shows the CNVs associated with complex genomic regions (i.e., within 5 kb of their breakpoints), including tandem repeats (TR), segmental duplications (SD), and pericentromeric regions (Peri). These CNVs accounted for 85.4% of all CNVs. The upper bar plot indicates the number of CNVs in each category or category overlap, and connected dots below represent the corresponding category combinations. (**E**) Phylogenetic clustering and a pairwise similarity heatmap based upon CNV alleles present in 40 inbred mouse strains. The clustering pattern was consistent with the known strain phylogeny ^61,62^: the four wild-derived strains (MOLF, CAST, SPRET, WSB), DBA1 and DBA2, and the NZW, NZB, and NZO strains formed separate clusters.

To investigate the advantage of using the T2T C57BL/6J backbone sequence, we examined the 225 protein coding genes (within the 208 Mb of sequence) in the T2T sequence that were absent in the GRCm39 reference sequence ^48^. We found that 131 CNVs could not be converted from their T2T coordinates to GRCm39 using LiftOver, indicating that these variants reside within genomic regions that are either absent, highly divergent, or structurally unresolved in the GRCm39 reference assembly (**Supplemental Data File S2**). This demonstrates that the use of the T2T reference sequence enables the identification of additional CNVs, which are present in complex genomic regions, that are incompletely represented using the conventional GRCm39 reference sequence.

We experimentally validated 10 of the identified deletion CNVs. For each CNV, the primers flanking the deleted region could only generate an amplicon from strains with the deletion allele (due to deletion-induced proximity of the primers), but not from C57BL/6J tissue (due to the large distance between the primers), which did not have the deletion. In contrast, PCR amplicons generated using primers within the deletion could only be amplified from C57BL/6J tissue, but not from strains with the CNV deletion allele (**Fig. S3** and **Table S2**).

### Gene-level CNV effects

The high-confidence CNVs were intersected with GENCODE-annotated protein-coding gene models, which led to the identification of 384 non-redundant CNV-affected protein-coding genes. Gene loss of function (LOF)-related effects predominated, which included 123 whole-gene losses, 132 exonic losses and 120 intronic losses. Together, LOF effects impacted 375 genes (97.7%). Gene gain-related effects were rare; only nine genes were assigned to the whole-gene gain, exonic gain or intronic gain classes (2.3%) (**Fig. 3A** and **Supplemental Data File 3**).

**Figure 3.**
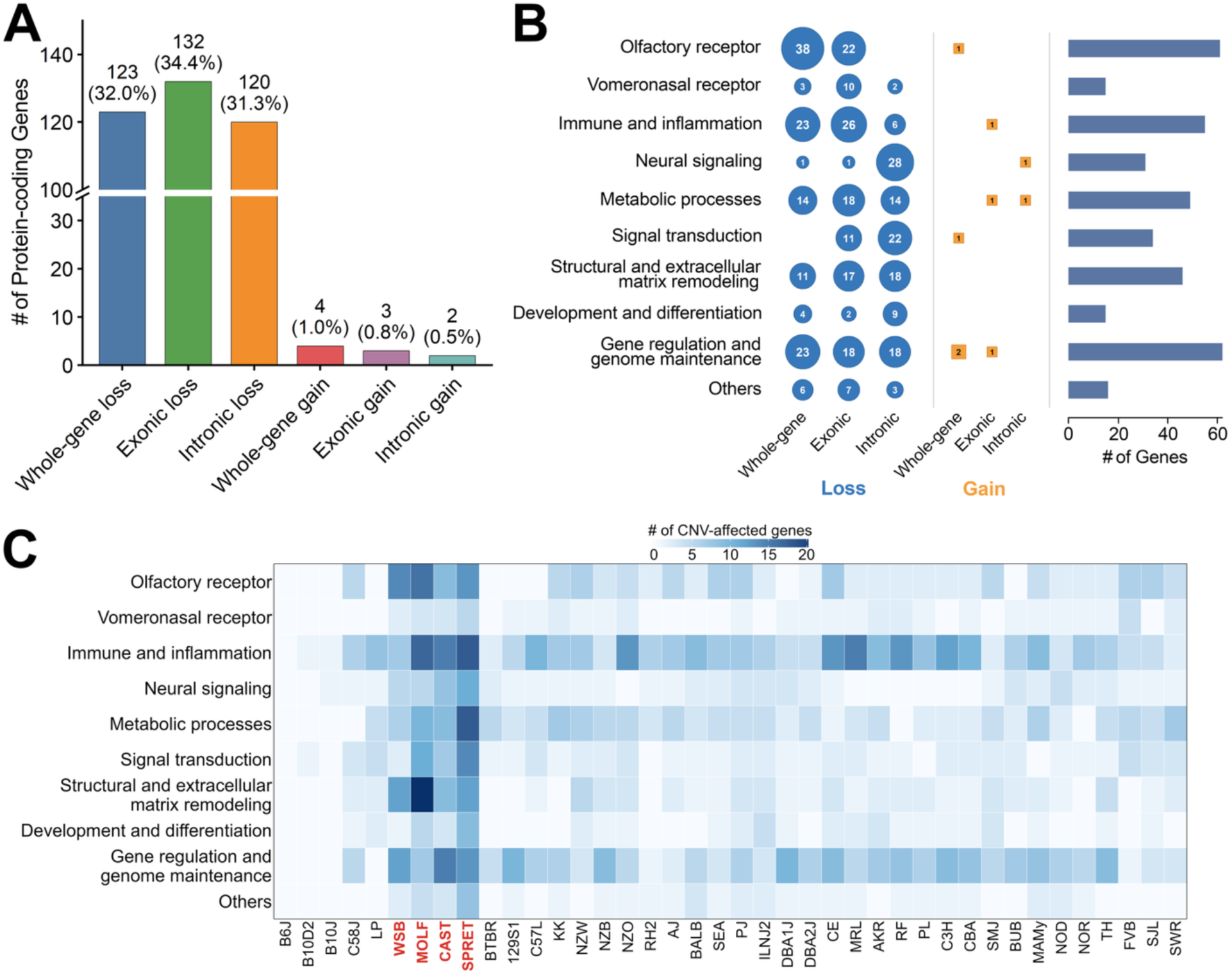
Functional annotation of CNV-affected protein-coding genes among the 40 inbred mouse strains. (**A**) A bar plot showing the number of non-redundant protein-coding genes that were assigned to six different classes of CNV effects: whole-gene loss, exonic loss, intronic loss, whole-gene gain, exonic gain and intronic gain. The numbers above bars indicate gene counts and percentages. (**B**) Functional classification of CNV-affected protein-coding genes across ten categories. The *left and middle panels* show the number of genes in each functional category affected by loss or gain events, respectively. The blue circles represent loss classes, whereas orange squares represent gain classes. The symbol size is proportional to the gene count, with the gene counts displayed inside each symbol. The *right panel* shows the total number of CNV-affected protein-coding genes in each functional category. (**C**) A heatmap showing the number of unique protein-coding genes affected by CNVs across functional categories and strains. The functional categories are shown in the rows and the 40 strains are in columns with the strain order matching the phylogenetic-tree order in Figure 2E. Color intensity indicates the gene count.

Functional classification revealed that the CNV-affected genes were distributed across diverse biological categories, with gene regulation and genome maintenance (62 genes), olfactory receptor (61 genes), immune and inflammation (56 genes), metabolic processes (48 genes), and structural and extracellular matrix remodeling (46 genes) representing the largest groups. Gene loss effects occurred across a wide group of functional categories; whereas gain effects were sparse, with no single category containing more than three gain-affected genes (**Fig. 3B**). A strain-level heatmap further revealed that there was a marked variation in the number and in the functional composition of CNV-affected genes across the strains, and the wild-derived strains (SPRET, MOLF, CAST and WSB) had the highest overall number of genes affected across functional categories (**Fig. 3C**).

### Pangenome expansion and sequence diversity

To characterize the extent of pangenome expansion across the 40 inbred mouse strains, we performed ordered-growth analysis using Panacus ^63^ on the Minigraph-Cactus pangenome graph ^55^. The cumulative pangenome size increased from 2.29 Gb for the initial strain (129S1) to 3.32 Gb after incorporation of all strains, representing an expansion of approximately 1 Gb beyond the baseline assembly (**Fig. 4A,C**). The amount of additional sequence contributed by each strain decreased progressively as more genomes were incorporated into the graph. Most classical laboratory strains contributed relatively modest amounts of additional sequence, whereas wild-derived strains introduced substantially larger amounts of novel sequence. CAST, MOLF, SPRET, and WSB contributed 99.98 Mb, 58.59 Mb, 225.47 Mb, and 25.12 Mb of additional sequence, respectively, which significantly exceeded the contributions of most of the classical strains (two-sided Wilcoxon rank-sum test, *P* = 2.8 × 10⁻³; **Fig. 4A**). Consistent with these observations, the pangenome growth curve did not reach a clear plateau after inclusion of all 40 strains, which indicates that additional sequence diversity remains to be captured from genetically divergent mouse populations (**Fig. 4C**).

**Figure 4.**
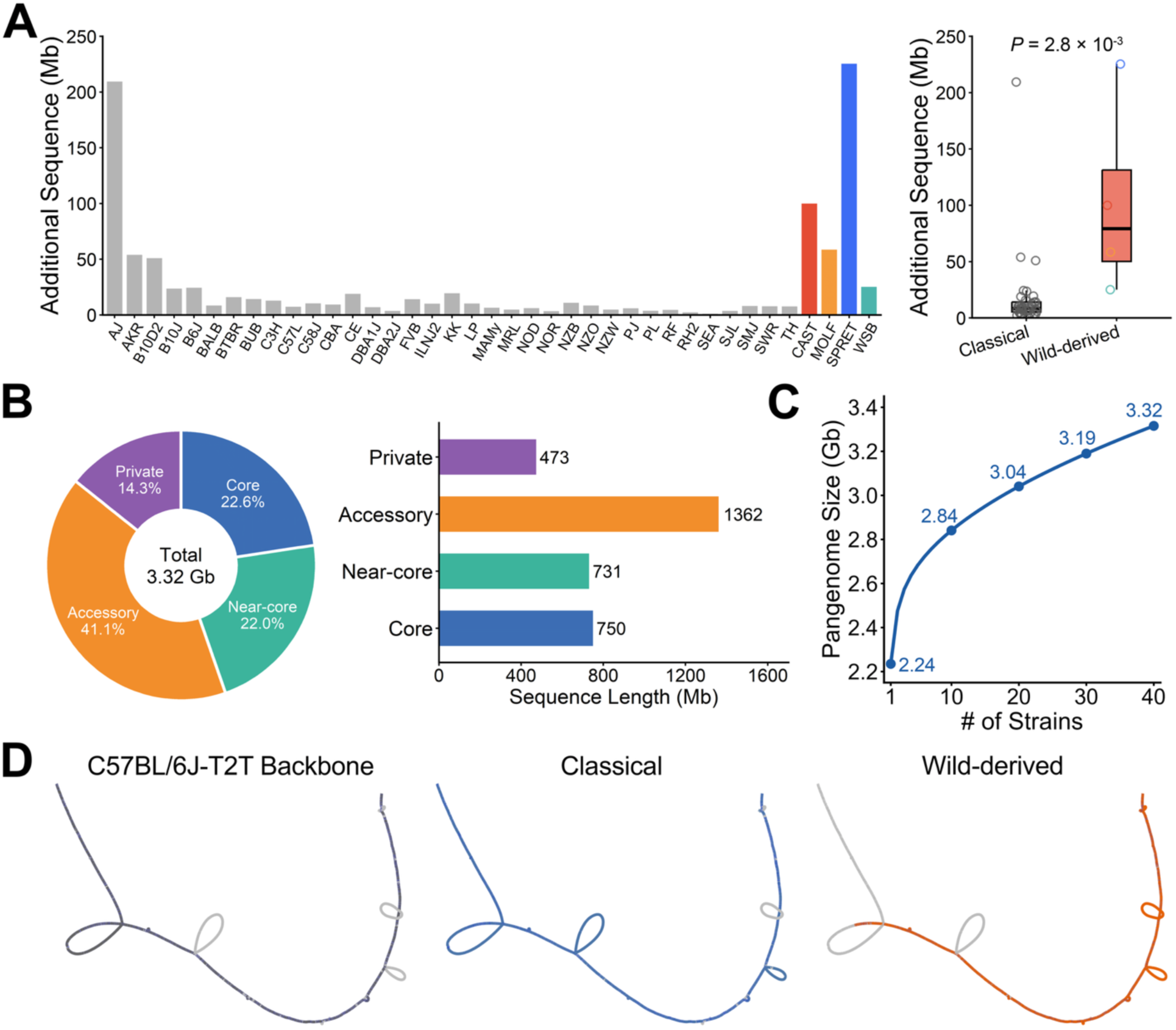
Mouse pangenome architecture and sequence diversity across 40 inbred strains. (**A**) Strain-specific contributions to mouse pangenome expansion were estimated using Panacus ^63^. Bars represent the amount of additional sequence incorporated into the pangenome following the sequential addition of each strain according to a predefined growth order consisting of 36 classical inbred strains followed by four wild-derived strains (CAST, MOLF, SPRET, and WSB). Because the analysis is order-dependent, additional sequence values reflect the increase in cumulative pangenome size at each growth step rather than the total amount of unique sequence present in each strain. The baseline strain (129S1), which served as the initial pangenome assembly, was excluded from the contribution analysis. The boxplot plot compares the additional sequence contributions between classical and wild-derived strains. Wild-derived strains contributed significantly more novel sequence to the mouse pangenome than classical laboratory strains (two-sided Wilcoxon rank-sum test, *P* = 2.8 × 10⁻³). (**B**) Composition of the mouse pangenome based on sequence-sharing frequency across the 40 strains. Sequences were classified as private (present in a single strain), accessory (present in 2–37 strains), near-core (present in 38–39 strains), or core (present in all 40 strains). The donut chart summarizes the relative proportions of each category within the pangenome, whereas the horizontal bar plot shows their corresponding sequence lengths. Core and near-core sequences accounted for 22.6% and 22.0% of the pangenome, respectively, while accessory and private sequences accounted for 41.1% and 14.3%. (**C**) The growth curve of the mouse pangenome. The cumulative pangenome size increased from 2.24 Gb to 3.32 Gb as the number of incorporated strains increased from 1 to 40, which indicates there is substantial sequence diversity across the strain panel and continued pangenome expansion with the inclusion of additional genomes. (**D**) Local pangenome graph visualization of a CNV hotspot spanning a 50-kb region on chromosome 11 (C57BL/6J-T2T coordinates: chr11:125,705,000–125,755,000), generated using BandageNG ^79^. *Left panel*: the C57BL/6J-T2T backbone path (dark gray). The *middle and right panels* highlight the paths among classical inbred (blue) or wild-derived (orange) strains. Multiple alternative graph branches and bubble structures reflect the CNV-associated sequence variation within this hotspot, which illustrates local sequence diversity among mouse strains. Distinct path usage patterns between classical and wild-derived strains indicate lineage-specific structural variation and demonstrate how CNVs contribute to local graph complexity.

Analysis of sequence-sharing patterns revealed a heterogeneous pangenome architecture (**Fig. 4B**). The core sequences present in all 40 strains accounted for 22.6% of the pangenome, while near-core sequences present in 38–39 strains contributed an additional 22.0%. Accessory sequences shared by a subset of strains represented the largest component of the pangenome (41.1%), whereas private sequences that were detected in only a single strain accounted for 14.3%. Together, these results demonstrate that a substantial fraction of the mouse pangenome consists of accessory and private sequences, highlighting the extensive genomic diversity across the classical and wild-derived mouse strains.

To investigate how CNVs shape the local pangenome architecture, we examined a highly variable CNV hotspot on chromosome 11 using graph-based visualization (**Fig. 4D**). This region displayed a complex network of alternative sequence paths, indicating extensive CNV-associated sequence diversity among the 40 mouse strains. Although they shared a common backbone structure, multiple lineage-specific path configurations were evident, which reflects differences in local copy-number state and sequence content. Numerous path bifurcations and rejoining events were observed throughout the graph, which illustrates how CNVs generate local graph complexity within the mouse pangenome. These observations illustrate how CNVs contribute to local pangenome complexity and sequence diversity among mouse strains.

### A CNV that contributes to the KK metabolic syndrome

We previously used a mouse genetic AI ^64^ to analyze an SV database and identify a candidate gene for mouse lymphoma ^9^. Therefore, this AI was used to analyze the CNV database and identify KK CNVs that could contribute to its metabolic syndrome. The AI analyzed the CNV alleles in KK and in 34 other strains that did not develop the metabolic syndrome, with the goal of identifying CNVs with KK alleles within genes associated with the metabolic syndrome (**Fig. 5A**). The 7 genes with a high impact CNV identified by the mouse genetic AI were then analyzed by a second AI. Gemini Pro DeepThink utilizes test-time compute scaling ^65,66^ and parallel thinking techniques to execute clinical grade differential logic across divergent biomedical areas, which surpasses that of earlier AIs. Gemini Pro DeepThink identified the high impact KK CNV in *NLR family pyrin domain containing 1B (Nlrp1b),* within a gene that was strongly associated with the metabolic syndrome as the likely causative candidate. This 29 kb deletion CNV located at chr11:81,817,572–81,846,589 in the T2T assembly (corresponding to chr11:71,057,660–71,086,677 in GRCm39) was only present in KK and NZW mice, and not in any of the 34 other analyzed strains. It deleted exons 2 through 6 of the 14 exons in the (canonical) *Nlrp1b* transcript (ENSMUST00000108515.9) (**Figs. 5B, S4**). RT-PCR analysis confirmed that full length *Nlrp1b* mRNA is expressed in the bone marrow of BALB/c mice, but not in KK mice. Consistent with the effect of the 29-kb CNV deletion that removes exons 2–6 of *Nlrp1b*, the KK transcript is markedly shorter because exon 1 is adjacent to exon 7 in the KK transcript (**Fig. S5**). *Nlrp1b* encodes an intracellular pattern recognition receptor that acts as an immune and metabolic sensor. In response to specific danger signals or intracellular stress, it forms a multiprotein complex (the NLRP1 inflammasome) that activates procaspase to caspase-1; which initiates a rapid inflammatory form of programmed cell death (pyroptosis) ^67,68^. Although inflammasomes drive inflammation, a functional *Nlrp1b* inflammasome plays a protective role in adipose tissue where it negatively modulates obesity-induced inflammation ^68^. *Nlrp1* deficient mice develop a metabolic syndrome ^69^ and transgenic expression of a functional *Nlrp1b* allele promoted lipolysis and protected against diet-induced insulin resistance ^70^. Since analysis of genetically engineered mice demonstrated that *Nlrp1b* impacts whether a metabolic syndrome will develop, we investigated whether the CNV-induced *Nlrp1b* deletion in KK mice altered two Nlrp1b-dependent cellular responses ^70^. Perilipin-1 associates with the surface of lipid droplets and protects them from lipolysis; and perilipin-1 levels in adipose tissue decrease when Nlrp1 activity is absent. Procaspase-1 is activated by the Nlrp1 inflammasome, and activated caspase 1 protects against developing the metabolic syndrome. Hence, if KK mice lack Nlrp1, it would be expected that perilipin-1 levels would decrease and procaspase-1 would increase (due to a failure to activate procaspase) in KK adipose tissue. Immunoblotting revealed that perlipin-1 was present in adipose tissue obtained from BALB/c mice, which has a very active *Nlrp1b allele*; but was not present in KK adipose tissue. In contrast, procaspase-1 was abundant in KK adipose tissue, but was only faintly detected in BALB/c adipose tissue (**Fig. 5C**). The changes in procaspase-1 and perilipin-1 levels in KK adipose tissue are consistent with the known downstream effects of a *Nlrp1b* deletion.

**Figure 5.**
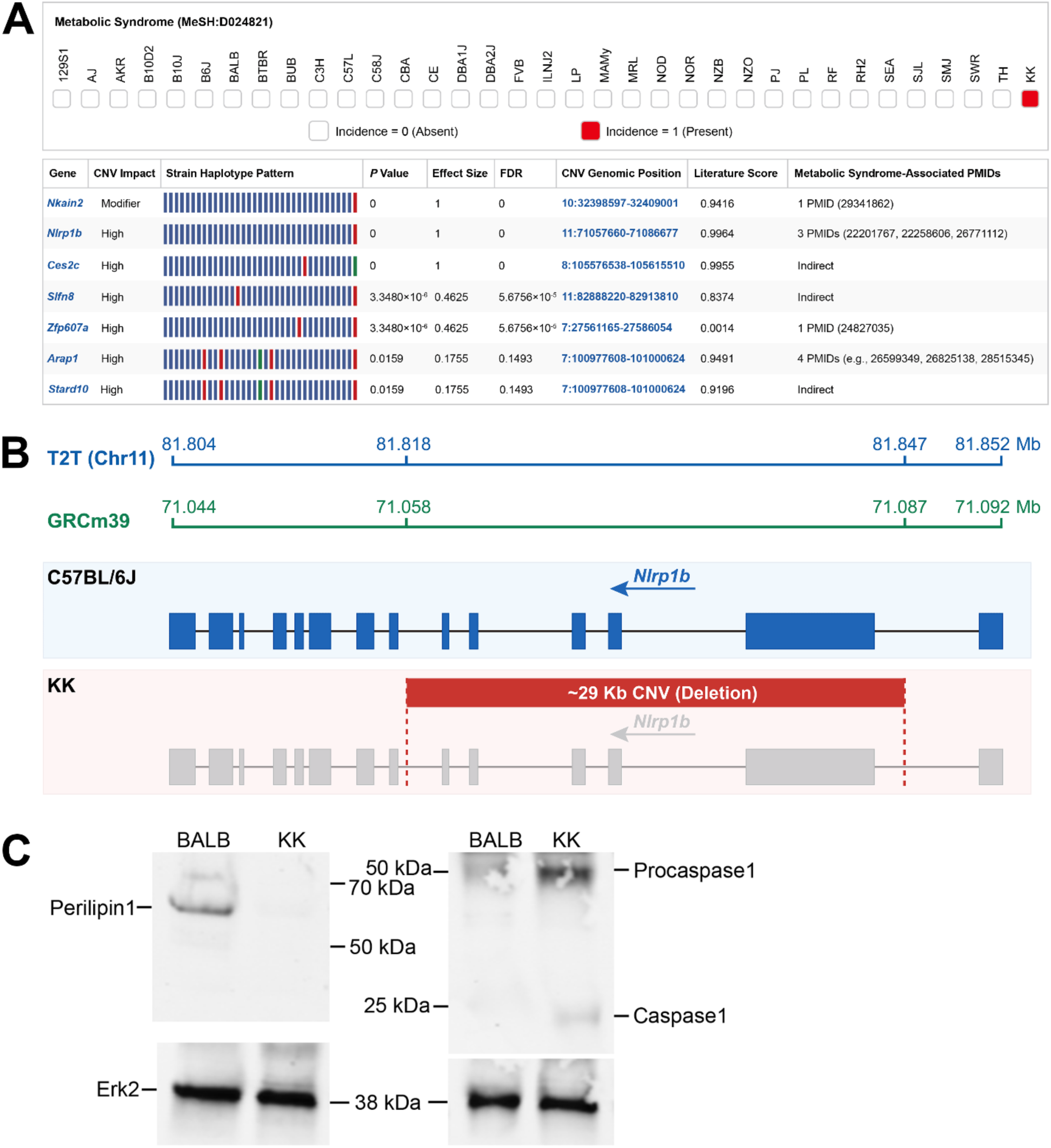
The AI pipeline identifies a genetic factor for the metabolic syndrome in KK mice. (**A**) *Top panel:* Metabolic syndrome was treated as a qualitative trait that appears in KK mice (incidence = 1), and not in the 34 other classical inbred strains (incidence = 0). *Bottom panel:* The AI pipeline performed a GWAS involving 35 strains to identify haplotype blocks with CNV allelic patterns that were only was only present in KK mice (i.e., genetic effect size = 1 and genetic association p-value = 0) and in genes that were associated with the metabolic syndrome. Two genes (indicated by gene symbol) with alleles that were only present in KK mice are shown at the top. Within the haplotype box: each block color represents a haplotype for one strain, strains with the same haplotype have the same color, and the blocks are shown in the same strain order as in the top panel. The chromosome and the starting and ending position of each CNV are also shown. The Literature Score represents the strength of the association of each gene with metabolic syndrome as determined by the AI-mediated literature search (MeSH term: Metabolic Syndrome D024821). PubMed identification numbers are provided for genes that have a direct link with the MeSH term. Otherwise, the AI indicates that the gene has an indirect association, which could result from interactions with other proteins that are associated with the metabolic syndrome. The CNV impact is determined by VEP analysis. Only *Nlrp1b* has a high impact CNV deletion that is directly associated with metabolic syndrome. *Nkain2* has an intronic CNV; *Ces2c* is indirectly associated with the metabolic syndrome; and the CNVs in *SLFn8, Zfp607a, and Arap1* are shared with one or more other strains. (**B**) Structural comparison of the *Nlrp1b* locus between C57BL/6J and KK mice. *Top panel*: The genomic coordinates of the *Nlrp1b* locus in the T2T assembly (chr11:81,803,840–81,852,194) and the GRCm39 reference genome (chr11:71,043,928–71,092,282) are shown. *Bottom panel*: Exon–intron structures of *Nlrp1b* are displayed for the C57BL/6J and KK strains. Arrowheads indicate the negative-strand orientation of *Nlrp1b*. The intact C57BL/6J *Nlrp1b* gene is shown in blue and serves as the reference for comparison with the KK strain. In KK mice, a 29,018 bp deletion CNV is highlighted by the red rectangle, and the two CNV breakpoints indicated by red dashed lines. This deletion removes exons 2 through 6 of *Nlrp1b*. The exons in the *Nlrp1b* gene that are disrupted by the KK CNV are shown in gray. (**C**) Analysis of perilipin-1 and caspase 1 expression in KK adipose tissue is consistent with the presence of a *Nlrp1b* deletion. Perilipin-1 and caspase 1 expression in mesenteric adipose tissues obtained from 9-week-old female BALB/c (body weight, 18 g) and yellow KK (body weight, 40 g) mice was examined by immunoblotting. Thirty ug of protein from each of the tissue samples were resolved on a 4-20% SDS-polyacrylamide gel, transferred to a nitrocellulose membrane, and immunoblotted with rabbit monoclonal anti-perilipin-1 or mouse monoclonal anti-caspase 1 antibodies, or with mouse monoclonal anti-Erk2 antibody as a loading control. While the KK and BALB/c adipose tissues had equivalent amounts of Erk2; perilipin-1 was present in BALB/c but not in KK adipose tissue; and procaspase-1 was abundant in KK adipose tissue and was only faintly detected in BALB/c adipose tissue.

### Human NLRP1 alleles are associated with human metabolic syndrome phenotypes

We wanted to investigate whether genetic variants within the human homologue of mouse *Nlrp1b* were associated with metabolic syndrome features in humans. The mouse *Nlrp* locus has undergone a lineage-specific gene duplication to create 3 paralogs (*Nlrp1a*, *Nalp1b*, *Nalp1c*), but humans have a single *NLRP1* gene ^68,71^, which is as the human orthologue of murine *Nlrp1b*. *NLRP1* alleles (rs11651270 and rs2670660) have been linked to susceptibility to Type 1 Diabetes in a Chinese Han population ^72^, but have not been linked with metabolic syndrome traits. Therefore, we examined *NLRP1* allelic associations with metabolic syndrome phenotypes in two deeply phenotyped large human population databases: the UK Biobank-based AstraZeneca PheWAS Portal (UKB) ^73^ and FinnGen release 13 PheWeb (FinnGen) ^74^. In the UKB, a 21.66-kb intragenic *NLRP1* deletion was nominally associated with three metabolic syndrome–related phenotypes, but none of them reached genome-wide significance (**Table S3**). In FinnGen, two intronic *NRLP1* INDEL variants were associated with alterations in BMI and body weight at genome-wide significant levels (*P* = 4.98×10^-8^ to 3.68×10^-10^, **Table S3**). However, their similar effect estimates indicate that they may represent a shared association signal rather than independent effects. Moreover, two *NLRP1* missense variants (rs2301582, rs11651270) were strongly associated with IGF-1 levels in the UKB population (*P*=3×10^-12^ to 1.3×10^-8^); and showed nominal associations with BMI and weight in the FinnGen population (*P*=1.71×10^-5^ to 3.71 × 10^-5^). The FinnGen SNP associations were retained (though >*P*=5×10^-8^) because they were the same coding variants identified in the UKB population. Hence, they provided cross-dataset, cross-phenotype support, which do not require independent replication at genome-wide significant levels. Collectively, these results link multiple classes of human *NLRP1* variation to glycemic, lipid, anthropometric and endocrine-metabolic traits, which supports the association of the KK *Nlrp1b* CNV with its metabolic syndrome.

## Discussion

This study generated a comprehensive T2T and pangenome-based CNV database covering 40 inbred mouse strains. Prior studies established that CNVs were an important part of mouse genome diversity. However, their ability to comprehensively characterize CNVs was limited by the genome assemblies, sequencing technologies, and the analytical strategies used at that time ^22,25^. More recent studies using T2T and LRS data have shown that repeat-rich, segmentally duplicated, pericentromeric, and strain-specific genome regions contain critical components of mouse genome diversity ^40,48^. For CNV discovery, we used the C57BL/6J T2T assembly as the reference because it provides a curated T2T representation of a key inbred strain ^48^. We focused the CNV database on autosomes (chromosomes 1–19) to maintain a consistent diploid framework for cross-strain comparison; sex chromosomes, particularly the PAR and Y-linked regions, require a separate sex-aware analysis due to differences in ploidy, homology and repetitive structure. This design provides a consistent autosomal T2T-pangenome framework for CNV discovery and interpretation. Many of the high-confidence CNVs in our database occur in genomic regions that are challenging to identify using conventional reference sequence-based analyses, and a subset of them could not be found using GRCm39 coordinates. By anchoring CNV discovery to the C57BL/6J T2T reference within a pangenome framework, our approach extends CNV analysis beyond the limitations of the earlier reference-based resources. Thus, the value of this database is not simply that it expands the number of CNVs; it makes complex genomic regions more accessible for CNV discovery. This CNV database provides a solid foundation for identifying candidate genetic factors for many important models of biomedical traits and diseases.

We demonstrate CNV database utility by using stacked AI analyses to identify a deletion CNV in KK mice that contributes to their metabolic syndrome ^49–51^. A functional *Nlrp1b* inflammasome plays a uniquely protective role in adipose tissue by negatively modulating obesity-induced inflammation ^68–70^. Since the *Nlrp1b* deletion allele is also present in NZW mice and NZW is a lean and metabolically normal strain, the *Nlrp1b* deletion cannot be a monogenic driver of the KK metabolic syndrome ^70^. Two prior analyses of intercross populations identified KK genetic loci on other chromosomes (7 and 9 ^75^; and 1, 6, 9 and 17 ^76^) that contribute to obesity and T2D. However, since both studies intercrossed KK mice with a strain (C57BL/6J) with an active *Nlrp1b* allele ^70^, they could not detect the KK *Nlrp1b* deletion effect. Hence, other genetic factors also contribute to the KK metabolic syndrome. Also, the F1 progeny of NZW and NZB mice provide a spontaneous model for systemic lupus-like autoimmunity ^77,78^. The NZW *Nlrp1b* deletion allele could contribute to its susceptibility to autoimmune disease, but we did not focus on that phenotype here. Nevertheless, our results indicate that additional analyses using this stacked AI analysis method and this comprehensive CNV database could identify the missing heritability for many murine other genetic models of human diseases and that murine results can guide the search for causative human genetic factors for human diseases that resemble the mouse pathology.

## Abbreviations

AI: artificial intelligence;
CNV: copy number variant;
LRS: long read sequence;
PAV: presence/absence variation;
SV: structural variant;
T2D: Type 2 Diabetes;
T2T: telomere to telomere;
TR: tandem repeats;

## RESOURCE AVAILABILITY

Further information and requests for resources and reagents should be directed to and will be fulfilled by the lead contact.

### Materials availability

This study did not generate new unique reagents.

### Data and code availability

The long-read sequencing (LRS) data is publicly available and has been deposited in the NCBI BioProject database under accession number PRJNA1250604.

All software and analytical methods used in this study are publicly available, as listed in the key resources table.

Any additional information required to reanalyze the data reported in this paper is available from the lead contact upon request.

## ACKNOWLEDGMENTS

This work was supported by NIH awards (1R01DC021133 and 1 R24 OD035408) to G.P.

## AUTHOR CONTRIBUTIONS

Conceptualization, G.P., W.R.; methodology, W.R.; formal analysis, software, and validation, W.R., Z.C.; visualization, W.R.; writing—original draft, G.P.; writing—review & editing, W.R. and G.P.; funding acquisition, G.P.; supervision, G.P.

## DECLARATION OF INTERESTS

The authors declare no conflict of interest.

## Notes

### Competing Interest Statement

The authors have declared no competing interest.

