## Supplemental Methods, Figures, Tables for "AI Analysis of a Copy Number Variant Database Identifies a Genetic Factor for a Murine Model of the Metabolic Syndrome"

**Supplemental Information**

### 4. Methods

#### 4.1 Sequencing Data and Genome Resources

Genomic DNA was isolated from 36 classical and four wild-derived inbred strains (CAST/EiJ, MOLF/EiJ, SPRET/EiJ, and WSB/EiJ); and long-read sequencing was performed using the PacBio Revio platform with the HiFi sequencing as previously described [1]. For each strain, HiFi reads were generated by circular consensus sequencing (CCS) using standard PacBio protocols, yielding an average of approximately 30× genome coverage per strain (mean 88.5 GB per strain). Only HiFi reads passing PacBio CCS quality filters were retained for downstream analyses. Detailed information on the strains, strain abbreviations, and sequencing depth is provided in Table S1. All analyses were performed using the complete telomere-to-telomere C57BL/6J reference genome (C57BL/6J-T2T, v1) [2], which was consistently used for read alignment, de novo sequence assembly, and pangenome inference. The analyses were restricted to autosomal chromosomes (chr1–chr19).

#### 4.2 Definition and Scope of the CNV Analysis

CNVs were defined as genomic segments exhibiting changes in copy number relative to the reference assembly, which was consistent with the established definition of CNVs as dosage-altering deletions (DEL) or duplications (DUP) occurring within reference-defined intervals. The analyses were restricted to medium and large-scale events with a minimum length of 10 kb, which is a threshold commonly applied to ensure robust CNV detection and reliable breakpoint resolution in complex genomic regions [3-5]. Only unbalanced CNVs that directly alter reference copy number were considered, including deletions and duplications [6]. Sequence insertions representing novel DNA segments that were absent from the reference genome (C57BL/6J-T2T, v1) were excluded, because these events are typically classified as insertional structural variants rather than coordinate-based CNVs.

#### 4.3 Pangenome Graph-Based CNV Discovery

A pangenome graph was constructed using the Minigraph-Cactus pipeline (v3.0.1) to enable graph-based CNV discovery across all strains [7]. The complete C57BL/6J-T2T reference genome was designated as the reference backbone [2], and de novo assemblies generated for each strain were incorporated as individual haploid paths. Input assemblies consisted of primary contigs produced by HiFiasm (v0.25.0) [8,9]. Pangenome construction and graph refinement were performed using Cactus (v3.0.1) in a single-machine configuration [7,10], which produced a graph representation of homologous genomic segments across all strains as discrete graph blocks. Copy number states were quantified by enumerating the number of strain-specific paths traversing each graph block relative to the reference path. CNVs were defined as deviations in block-level copy number compared with the reference genome, with corresponds with deletions (reduced copy number) or duplications (increased copy number) of reference-defined intervals.

#### 4.4 Assembly-Based CNV Discovery

De novo genome assemblies were generated independently for each strain and used for CNV discovery. HiFi reads were assembled using HiFiasm in primary contig mode to produce haploid assemblies for downstream analysis [8]. Primary contigs from each strain were aligned to the C57BL/6J-T2T reference genome using minimap2 (v2.30) [11], with alignment presets selected according to expected sequence divergence between the assemblies and the reference sequence: asm5 for classical inbred strains and asm10 for wild-derived inbred strains.

Copy number altering events were identified from assembly-reference alignments using SVIM-asm (v1.0.3) in haploid mode [12], which directly reports deletions and duplications based on alignment discordance and split contigs. In parallel, PAV (v2.4.6) was used to identify deletions and insertions from haploid assemblies relative to the reference genome [13]. The insertions were subsequently evaluated using a custom post-processing pipeline to distinguish copy number gains from novel, non-reference sequence insertions. Only events corresponding to reference-defined deletions or duplications were retained for downstream CNV analyses.

#### 4.5 Breakpoint and Read-Depth Based CNV Discovery

PacBio HiFi reads from each strain were aligned to the C57BL/6J-T2T reference genome using minimap2 with presets optimized for PacBio HiFi sequencing data [11]. The alignments were sorted and indexed to generate coordinate-sorted BAM files for downstream analyses with Sawfish (v2.2.1) [14]. CNV discovery was conducted independently for each strain using the Sawfish discover workflow, which integrates breakpoint evidence from split and discordant alignments with read-depth information to identify candidate copy number changes. Regions prone to mapping artifacts or unreliable copy number estimation were excluded using a predefined CNV exclusion mask, and a minimum variant size of 50 bp and a minimum mapping quality threshold of 5 were applied. Per-sample discovery results were subsequently combined across all strains using the

Sawfish joint-call workflow to generate a unified multi-sample CNV callset, which enabled consistent genotyping and comparison of copy number states across the strains. From the Sawfish callset, only the copy number altering variant classes (DEL, DUP, and CNV records representing quantitative copy number changes) were retained.

#### 4.6 CNV Filtering and Exclusion Criteria

To ensure high-confidence CNV detection, a unified filtering framework was applied to CNV callsets generated by pangenome-based, assembly-based, and read-depth–based analyses prior to integration. Across all methods, only variants passing the internal caller-specific quality control filters (FILTER = PASS) were retained. Analyses were restricted to copy number altering variant classes, including deletions and duplications as defined by each caller, and to variants with an absolute length of ≥10 kb. Redundant variant representations were collapsed by retaining, for each genomic locus, the record(s) with the largest span and by removing strict duplicates with identical coordinates and lengths.

To eliminate genomic regions prone to mapping artifacts or unreliable copy number estimates, a comprehensive CNV exclusion mask was constructed for the C57BL/6J-T2T reference genome. The mask includes annotated assembly gaps, ribosomal RNA (rRNA) repeats, SYNREP and GSAT satellite repeats, and telomeric regions. Annotation tracks were obtained from the UCSC Genome Browser or the C57BL/6J-T2T assembly (GCA_964188535.1) [15], and individual BED tracks were combined, sorted, and merged to generate a non-redundant exclusion mask. CNVs whose genomic intervals overlapped the exclusion regions by ⩾50% were removed. Additional caller-specific filters were applied where appropriate. For CNVs derived from Sawfish, the variants were required to have QUAL ⩾ 30 and SVCLAIM values indicating adjacency-supported events (J) or combined depth and adjacency support (DJ), excluding depth-only calls. For CNV-type records reporting quantitative copy number estimates, at least one strain with a homozygous genotype and depth-based copy number quality scores (CNQ) ≥ 13 were required, whereas deletion and duplication calls required at least one homozygous strain. Variants that were homozygous in ⩾20 strains were excluded to remove strain-fixed or reference-specific events. For assembly-based CNVs, SVIM-asm calls were restricted to deletions and tandem duplications (DUP:TANDEM), while PAV-derived calls were restricted to deletions and insertions, with large insertions retained as candidate copy number gain signals for subsequent integration.

CNVs shared by >20 strains were removed because they are usually artifacts. The rational for this is that any CNV that appears as a homozygous alternate (1/1) in over half of the strains is unlikely to represent a segregating structural variant. Instead, they often result from systematic biases, such as reference misrepresentation (e.g., collapsed or expanded regions in the reference genome) or recurrent assembly/alignment artifacts occurring within complex genomic regions (e.g., segmental duplications or tandem repeats). These biases generate consistent signals across many strains, which produces "shared CNVs" that are technical artifacts rather than true variants.

**4.7 Imputation of Missing CNV Genotypes**

In the assembly-based CNV callset, which was derived from the integration of SVIM-asm and PAV identified CNVs, some of the CNVs lacked genotype assignments across strains. To address this, genotypes were imputed using three complementary approaches: cuteFC (v1.0.2) [16], Sniffles2 (v2.7.1) [17], and SVJedi-graph (v1.2.1) [18]. cuteFC and Sniffles2 leverage read alignment signals to infer breakpoint-supported genotypes, whereas SVJedi-graph performs genotyping within a graph-based framework. The genotype calls were integrated across the different methods, and a consensus genotype was assigned when at least two of the three approaches produced concordant results; the strains without such agreement were retained as missing in subsequent analyses. For CNVs identified through the pangenome graph-based Minigraph-Cactus pipeline, missing genotypes were imputed using the SVJedi-graph alone because of its compatibility with graph-based genome representations and its ability to directly exploit the pangenome structure for genotyping.

**4.8 Integration and Confidence Stratification of CNV Callsets**

Assembly-based CNV callsets generated by SVIM-asm and PAV were merged using Jasmine (v1.1.5) [19], clustering variants with a minimum reciprocal overlap of 50% (min_overlap = 0.5), a maximum breakpoint distance of 1 kb (max_dist = 1000), and a linear distance scaling factor of 0.1 (max_dist_linear = 0.1). Structural variants identified from the pangenome graph-based Minigraph-Cactus pipeline were consolidated using Truvari (v5.4.0) [20] in collapse mode, merging highly similar variants into representative events based on breakpoint proximity, reciprocal overlap, and size similarity. Variants were considered collapsible if breakpoints were within 1 kb, reciprocal overlap was ⩾70%, and size similarity was ⩾70%; all variants were retained irrespective of supporting sample count, and numeric INFO fields were summarized using median values. Finally, CNV callsets from the three complementary strategies (pangenome graph-based, assembly-based, and breakpoint/read-depth-based approaches) were integrated using SURVIVOR (v1.0.7) [21], merging variants within a 2 kb breakpoint distance while requiring consistent variant types. A minimum variant size of 10 kb was applied throughout to ensure consistency.

CNVs were stratified into four tiers with different confidence levels based on the number of supporting discovery strategies and the level of evidence provided by each method, which included pangenome graph-based detection, assembly-based detection, and breakpoint/read-depth-based detection. Within each method, CNVs supported by multiple independent evidence signals or tools (e.g., breakpoint and read-depth signals in Sawfish, or concordant detection by both PAV and SVIM-asm in the assembly-based approach) were classified as high confidence (HC), whereas those supported by a single evidence source were classified as moderate confidence (MC). Variants supported by at least two independent methods, including at least one HC source, were classified as Tier 1a (strong orthogonal support), while those supported by at least two methods, but lacking HC support were assigned to Tier 1b (orthogonal support). Variants identified by a single method with HC evidence were classified as Tier 2 (strong single-source support), whereas those supported by a single method with only MC evidence were designated as Tier 3 (candidate variants).

#### 4.9 Segmental Duplication Annotation

Gene-level segmental duplications were annotated independently for each of the 40 strains using SegDupAnnotation2 (v1.3.3) [22]. For each strain, haplotype-resolved assemblies generated by Hifiasm were used as input together with the GENCODE vM38 protein-coding transcript set as the reference gene model. Long-read HiFi sequencing data were incorporated to support duplication resolution and copy number inference. Gene models were required to achieve at least 90% alignment coverage to be retained (min_gene_model_alignment = 0.90), ensuring high-confidence mapping of annotated transcripts to duplicated regions. Uncharacterized loci were excluded based on gene name prefixes (e.g., “LOC”) to focus analyses on well-annotated genes, and haploid chromosomes were automatically detected to avoid inflation of copy number estimates. Default workflow parameters were otherwise applied. The resulting annotations were used to quantify gene duplication events and to support downstream analyses of segmental duplication patterns across strains.

#### 4.10 Distribution and Sharing Patterns of CNVs

A total of 1,594 high-confidence (T1a, T1b, and T2) CNVs were used for downstream analyses. The number of CNVs identified in each of the 40 mouse inbred strains was calculated based on the presence of CNV alleles in individual strains. CNV allele sharing patterns were evaluated by determining the number of strains harboring each CNV allele across the population. To compare CNV size distributions between classical and wild-derived mouse strains, CNVs identified in the 36 classical inbred strains and four wild-derived strains (MOLF, CAST, SPRET, and WSB) were separately collected. CNV lengths were calculated based on genomic span, and statistical significance between the two groups was assessed using a two-sided Wilcoxon rank-sum test.

Tandem repeat (TR) annotations were obtained from our previously established tandem repeat resource, which were generated using the same mouse strains [23]. Segmental duplication (SD) annotations were derived from the analyses described in Section 4.9. Pericentromeric region (Peri) annotations were downloaded from the UCSC Genome Browser based on the GCA_964188535.1 assembly. CNVs were considered associated with TRs, SDs, or pericentromeric regions if 5-kb breakpoint-flanking regions overlapped the corresponding annotations. To investigate strain relationships based on CNV variation, a binary CNV genotype matrix was generated, where rows represented mouse strains and columns represented CNVs. Presence and absence of CNV alleles were encoded as 1 and 0, respectively. Hierarchical clustering and pairwise similarity heatmap visualization were performed using the R package ComplexHeatmap.

**4.11 Functional Annotation of CNV-affected Protein-coding Genes**

CNV-affected protein-coding genes were identified by intersecting the 1,594 high-confidence CNVs with GENCODE vM38 gene annotations, and this identified CNV effects in 384 non-redundant genes. Based on the affected gene-body region and CNV type, deletion CNVs were classified as: whole-gene loss, exonic loss or intronic loss. Duplication CNVs were classified as whole-gene gain, exonic gain or intronic gain. For genes associated with multiple effects, a single representative class was retained using the hierarchy: whole-gene > exonic > intronic.

The classification of the functional effect of a CNV was primarily based on UniProtKB protein annotations obtained by mapping gene names to UniProt entries restricted to *Mus musculus* [24], with MGI Mammalian Phenotype annotations [25,26] used as supporting evidence. Each gene was manually assigned to one primary category: olfactory receptor, vomeronasal receptor, immune and inflammation, neural signaling, metabolic processes, signal transduction, structural and extracellular matrix remodeling, development and differentiation, gene regulation and genome maintenance, along with a few other categories. Unique gene counts were summarized across CNV-effect classes and functional categories. A strain-level heatmap was generated to show the number of CNV-affected unique protein-coding genes in each functional category across the 40 strains.

#### 4.12 Pangenome Analysis and Graph Visualization

Pangenome analyses were performed on the Minigraph-Cactus graph constructed from the 40 inbred mouse strains. Panacus (v0.4.2) [27] was used to quantify pangenome growth and sequence-sharing patterns across strains. Ordered-growth analysis was performed to estimate cumulative pangenome expansion and the incremental sequence contribution of each strain, whereas histogram-based analysis were used to classify graph sequences into core, near-core, accessory, and private components according to their sharing frequencies across the strain panel. To investigate local graph structure, a highly variable CNV hotspot on chromosome 11 (C57BL/6J-T2T coordinates: chr11:125,705,000–125,755,000) was extracted from the pangenome graph using ODGI (v0.9.4) [28], optimized, and converted to GFA format. The resulting subgraph was visualized with BandageNG (v2026.6.1) [29] to examine local graph topology, including alternative paths and bubble structures associated with CNV diversity among mouse strains.

#### 4.13 Experimental Validation of Deletion CNVs

Mouse genomic DNA was prepared from liver tissues (~20 mg) obtained from 129S1, AJ, AKR, BTBR, CBA, DBA2J, KK, LP, NOD, NZB, SJL, TH and B6J mice using the PacBio Nanobind tissue kit. Liver tissue from C58J and PJ mice were gifts from the Jackson Laboratory. The GoTaq Green Master Mix was used to amplify DNA fragments in or near the sequences where CNV deletions were identified. For each CNV deletion, a pair of primers flanking the deleted region were used (Table S2, primers ending with F and R), which could only produce an amplicon from tissue obtained from mice with the deletion. C57BL/6J genomic DNA had the deleted region for all the CNVs. Hence an amplicon from C57BL/6J tissue could not be generated because it was too long. For controls, two internal primers were designed to pair with the flanking primers (i.e., primers ending with R1 are paired with F, and F1 is paired with R). When one of the primers is an internal primer, a fragment can be amplified from C57BL/6J tissue, but not from tissue obtained from strains with the deletion.

#### 4.14 Analysis of *Nlrp1b* mRNA

Total RNAs were purified from the bone marrow of the long bones (femurs, tibias, humeri) of Balb/c (Ba) and KK-Ay (KK) mice using the TRIzol Reagent (Thermo Fisher). One ug each of total RNA were reverse transcribed using the High-Capacity cDNA Reverse Transcription kit (Applied Biosystems) in a 20-µl volume. One µl each of the cDNAs were then subjected to PCR amplification using the GoTaq® G2 DNA polymerase master mix (Promega) and primers for *Nlrp1b* or *Gapdh* (control) transcripts. RT-PCR amplicons of the bands indicated in Fig. S5 were sequenced by Sanger sequencing (McLab, South San Francisco) when only one major amplicon was present on the gels.

*Nlrp1b* and *Gapdh* transcript primers: To amplify the cDNA of the *Nlrp1b* transcript from Balb/c and KK-Ay mice, the cDNA sequences of KK-Ay mice and Blab/c were aligned before the primers were designed. The *Nlrp1b* cDNA sequence was inferred from the KK LRS data and compared to the *Nlrp1b* GRCm39/mm39 reference sequence. The *Nlrp1b* cDNA of Balb/c is available in GenBank (GenBank: DQ117583.1). *Nalp1b* primers used: (1) N1b-F: ACAGTATCTGAGGGCTCTGTACCA (for KK-Ay mice in the "Nlrp1b, FR1" reaction) and N1b-F_Ba: GCAGTATCTGAGGGCTCTATCATCAT (for Balb/c mice in the "Nlrp1b, FR1" reaction). Both N1b-F and N1b-F_Ba target the first exons of the *Nlrp1b* transcripts. Since the 29 kb-genomic deletion in KK-Ay mice encompasses exons 2 to 6, it is necessary to use forward primers targeting the first exon to see a difference between KK-Ay and the Balb/c control. (2) N1b-F1: CAAGGATGGTTCTGTACAGGTGGA. (3) N1b-R1: GGCAATTGAACTCGGTACAGGTT. (4) N1b-R: AGGAAAAGAGATGCCAGGCTCCTAT. (5) *Gapdh* transcript primers: Gapdh-F2: GTAGACAAAATGGTGAAGGTCGGT and Gapdh-R1, GGTCCAGGGTTTCTTACTCCTTG were used for *Gapdh* transcript amplification from KK-Ay and Balb/c tissue.

**4.15 Perilipin-1 and Caspase 1 Expression in Adipose Tissue**

Mesenteric adipose tissues were obtained from 9-week-old female BALB/c (body weight, 18g) and yellow KK (body weight, 40 g) mice and were homogenized in RIPA buffer supplemented with a protease inhibitor cocktail (Sigma P8340, 1 to 100) using a Precellys tissue homogenizer. Thirty ug of protein from each sample were resolved on a 4-20% SDS-polyacrylamide gel and transferred to a nitrocellulose membrane and immunoblotted with the following antibodies: rabbit monoclonal anti-perilipin-1 antibody (Cell Signaling, #9349T, 1:1000 dilution) or mouse monoclonal anti-caspase 1 antibody (Santa Cruz Biotechnology, sc-56036, 1:1000 dilution) or mouse monoclonal anti-Erk2 antibody (Santa Cruz Biotechnology, sc-1647, 1:5000). IRDye® 680LT goat anti-rabbit IgG (925-68021) or IRDye® 800CW goat anti-mouse IgG (Licor, 925-

32210) were used as the secondary antibodies. After blotting, the membrane was scanned using a Licor Odyssey imaging system.

#### 4.16 Human *NLRP1* Variant Associations with Metabolic Traits

To assess whether alleles in the human homologue of murine *Nlrp1b* affected susceptibility to features of the metabolic syndrome, publicly available association statistics for the human *NLRP1 l*ocus (Ensembl gene ID: ENSG00000091592) were examined using data obtained from the UK Biobank–based AstraZeneca PheWAS Portal [30] and FinnGen release 13 PheWeb [31]. CNVs, INDELs, and SNPs located entirely within the genomic boundaries of human *NLRP1* (GRCh38, chr17:5,499,415–5,619,424) were examined for associations with metabolic traits, which included glycemic levels, lipid levels, anthropometric measurements, and endocrine traits.

Variant class, identifiers, genomic positions, predicted annotation, phenotype, effect estimate, *P* values, and sample sizes were recorded as in the respective databases. CNV sizes were calculated using the reported GRCh38 breakpoint coordinates. Since the FinnGen BMI and weight traits were inverse-rank normalized, the effect estimates for these traits represent transformed trait units. The results were summarized descriptively without meta-analysis because the phenotype definitions, association models, and effect scales differed between the databases. No individual-level genotype or phenotype data were accessed or analyzed.

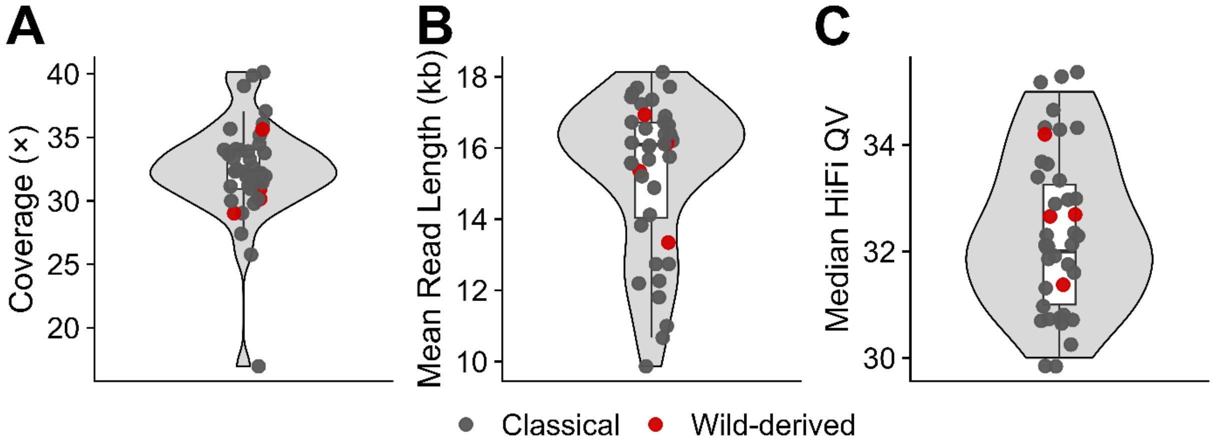


**Figure S1.** Summary of PacBio HiFi sequencing quality metrics across 40 inbred mouse strains. (**A**) The distribution of sequencing coverage across the 40 mouse strains. Except for SMJ, which exhibited the lowest sequencing depth (17×), most strains achieved coverage levels between 30- and 35-fold genome coverage. This provided sufficient depth for high-confidence genome assembly and copy number variant (CNV) discovery. (**B**) The distribution of mean HiFi read lengths across the 40 mouse strains. Most strains exhibited mean read lengths ranging from 16 to 18 kb, which is consistent with high-quality long-read sequencing. Three strains, C58J, RH2, and SMJ, showed comparatively shorter mean read lengths of approximately 10 kb. (**C**) The distribution of median HiFi read quality values (QV) across the 40 mouse strains. All strains achieved median HiFi QV greater than 30, corresponding to a base-level accuracy exceeding 99.9%, which indicates that there is consistently high sequencing quality across the dataset.

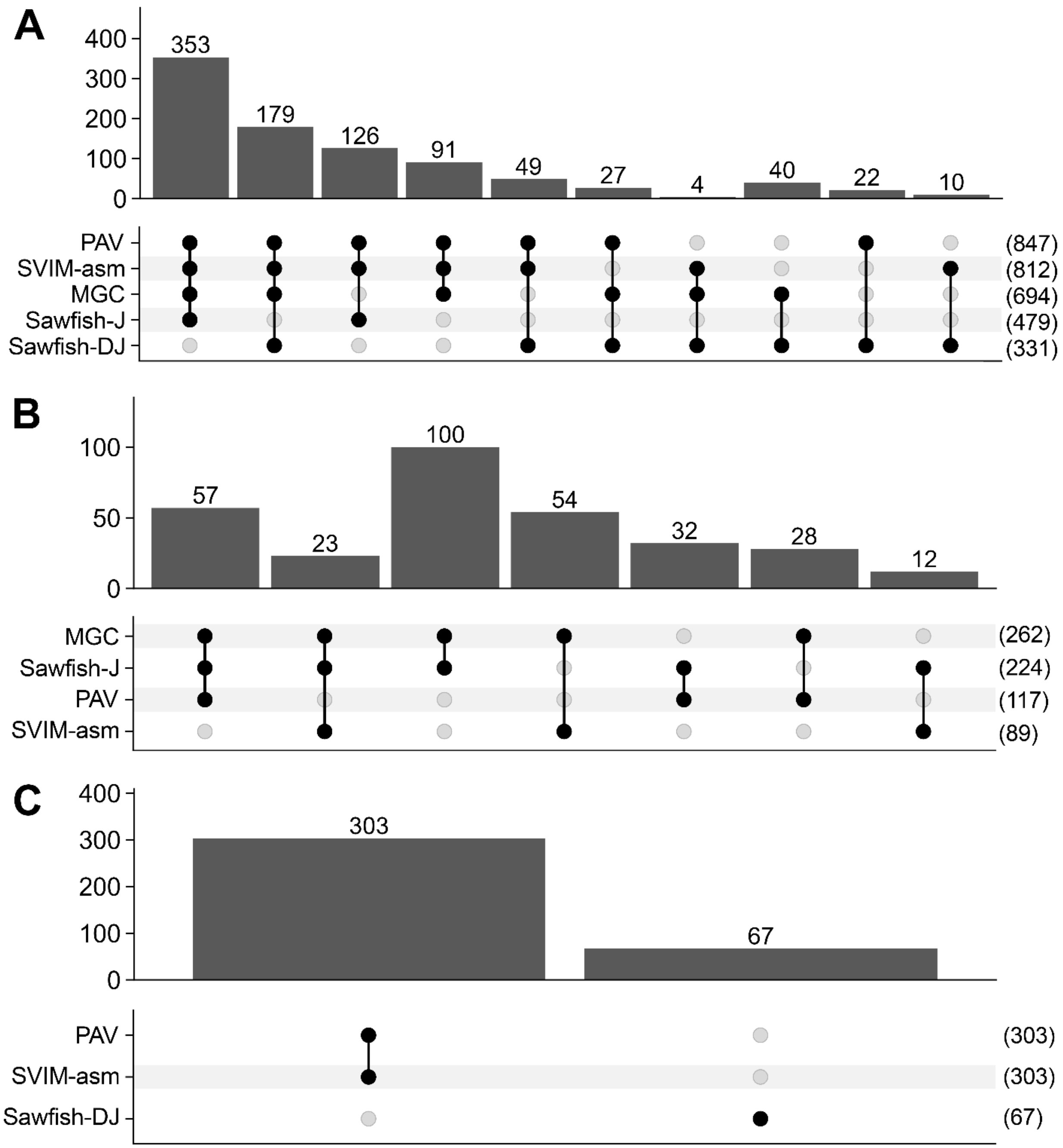


**Figure S2.** Concordance among deletion CNVs identified by the different detection methods. (**A**) UpSet plot showing the overlap of deletion CNVs (DELs) assigned to the T1a category across multiple detection methods. Minigraph-Cactus (MGC) represents the pangenome graph-based strategy. Sawfish-J corresponds to adjacency-supported events and was considered a moderate-confidence breakpoint/read-depth call, whereas Sawfish-DJ represents depth- and adjacency-supported events and was considered a high-confidence breakpoint/read-depth call. Assembly-based CNVs detected by both PAV and SVIM-asm were classified as high confidence, whereas CNVs detected by only one of the two methods were classified as moderate confidence. The numbers in parentheses indicate the total number of CNVs identified by each method. More than 90% of T1a CNVs were supported by at least three methods, including at least one high-confidence strategy. PAV identified the largest number of CNVs (n = 847). (**B**) UpSet plot showing the overlap of deletion CNVs assigned to the T1b category. These CNVs were supported by at least two independent discovery strategies but lacked support from any high-confidence strategy. Among all methods, Minigraph-Cactus identified the largest number of CNVs (n = 262). (**C**) UpSet plot showing the overlap of deletion CNVs assigned to the T2 category. These CNVs were supported exclusively by a single high-confidence strategy and therefore represent a confidence class that is distinct from T1a and T1b CNVs.

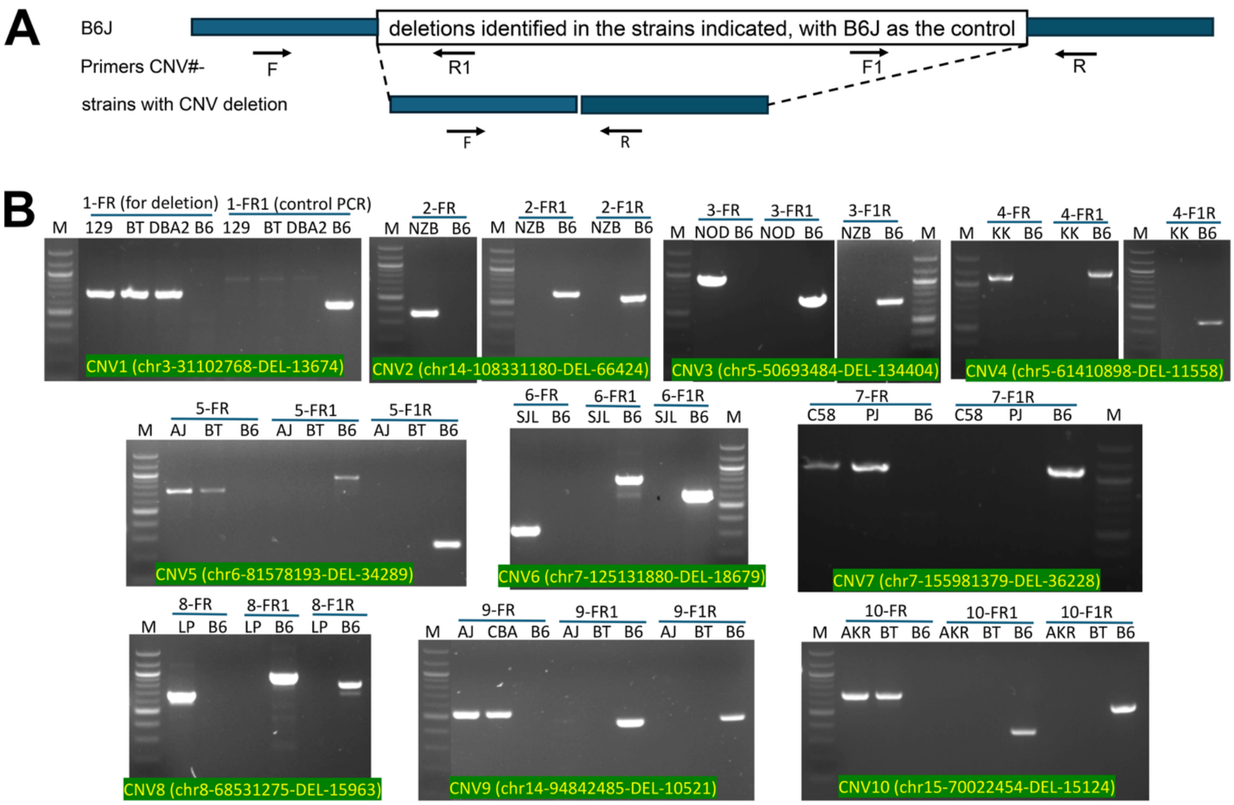


**Figure S3**. Experimental validation of 10 CNV deletion alleles by PCR amplification. (**A**) A schematic illustration of a region where a deletion CNV was identified by sequence analysis. The C57BL/6J (B6J) genome had the deleted region, which was absent in the other strains examined. The PCR primers ending with F and R are located far apart in the B6 genome, which is why these primers do not generate an amplicon when the PCR is performed using B6 tissue. For strains with CNV deletion alleles, PCR fragments of the expected sizes were amplified by these primers due to deletion-induced proximity. Primer pairs F and R1 or F1 and R only amplified fragments from B6 of the expected sizes shown in Table S2. (**B**) The gels showing the PCR amplicons generated by PCR analysis of tissue obtained from the indicated strains using the indicated primers. The CNV number and its chromosomal location are shown at the bottom of each gel. fragments. M: 100 bp ladder from New England Biolab.

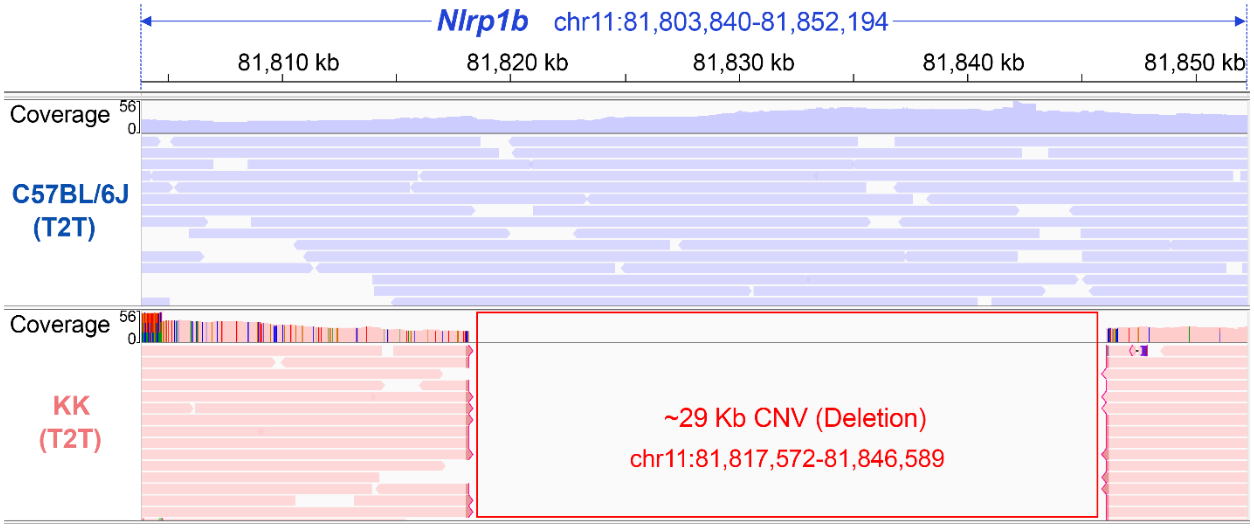


**Figure S4**. IGV visualization of PacBio HiFi read alignments from our C57BL/6J and KK LRS mapped to the C57BL/6J-T2T reference genome. Coverage and read-alignment tracks are displayed across the candidate CNV locus. The CNV interval, indicated by the red rectangle, is characterized by the near-complete absence of read coverage and aligned reads in KK relative to our C57BL/6J sequence, which provides direct LRS support for this deletion CNV.

**A**

```
Nlrp1b_mRNA_Ba GTCTCCTCTATCAG----TATCAAATACCCATTCCCAGA-AAGAAATTT-TGG-CAGAGA   53
Nlrp1b_mRNA_KK GTCTCCTCTATGAGTTCTGATTAAATACCCATTCCCAGAGCAGAAATTTCTGGCCA-TGA   59

Nlrp1b_mRNA_Ba CGAGAAGCCACTGGACTCCTACTTGGAACAACAGAAGAGTGTCTTGAAGTAATAAGAGCA  113
Nlrp1b_mRNA_KK CAAGAAGACACTGCTCTCCTGCTTGCAACATCAGAAGAGAGTCTTGAAGAAATAAGAGCA  119
                                N1b-F_Ba
Nlrp1b_mRNA_Ba CAATTTGCTCAGAGAAGCAGTATCTGAGGGCTCTATCATCATGGAAGAATCCCCACCCAA  173
Nlrp1b_mRNA_KK CAAACTGCTCAGAGAAACAGTATCTGAGGGCTCTGT-ACCATGGAAGAATCTCAGTACAA  178
                                 N1b-F
Nlrp1b_mRNA_Ba GCAGAAAAGTAACACAAAGGTTGCTCAGCATGAAGGTCAACAAGACTTGAACACAACGAG  233
Nlrp1b_mRNA_KK GCAGGAACATAACAAAAAGGTAGCTCAGGATGAAGGTCAAGAGGACAAGGACACCATCTT  238

Nlrp1b_mRNA_Ba ACATATGAATGTAGAGCTGAA-GCACAG--------------ACCCAAGCTAGAGAGACA  278
Nlrp1b_mRNA_KK CGAAACAATAGAAGCTATAGAGGCAAAGCTGATGGAGCTCAAAACCAACCCAGAGAGTAC  298

Nlrp1b_mRNA_Ba CTTGAAGCTAGGAATGATTCCAGTAGTATATATGAAGCAGGGAGAAGAGATACTTTACCC  338
Nlrp1b_mRNA_KK CTTCAATTATG-------------------------------------------------  309

---                                                             N1b-F1
Nlrp1b_mRNA_Ba GAGGACTCTTCAGCTGAATATGGAAAAACAGCAAGGATATGCACTGATATCCCCAAGGAT 1958
Nlrp1b_mRNA_KK ------------------------------------------------------------  309

Nlrp1b_mRNA_Ba GGTTCTGTACAGGTGGACCCCAATCACTAATGCCAGTTGGGAGATTCTCTTCTACAATCT 2018
Nlrp1b_mRNA_KK ------------------------------------------------------------  309

---

Nlrp1b_mRNA_Ba AGAAGATATCCTGACCTCATTCAAGCAGCAGAGACAACAGTCAGGAGCCAATCCCATGGA 2678
Nlrp1b_mRNA_KK --------------------------------------------GAGCCAATCCCATCGA  325

Nlrp1b_mRNA_Ba AATTCTGGGGACTGAAGAAGACTTCTGGGGCCCTATAGGACCTGTGGCTACTGAGGTGGT 2738
Nlrp1b_mRNA_KK AATTCTGGGGACTGAAGAAGACTTCTGGGGCCCTATAGGACCTGTGGCTACTGAGGTGGT  385
                              N1b-R1
Nlrp1b_mRNA_Ba TTACAGAGAAAGGAACCTGTACCGAGTTCAATTGCCCATGGCTGGTTCCTACCACTGTCC 2798
Nlrp1b_mRNA_KK TTACAGAGAAAGGAACCTGTACCGAGTTCAATTGCCCATGGCTGGTTCCTACCACTGTCC  445

Nlrp1b_mRNA_Ba CAGCACAAGACTCCACTTTGTAGTGACAAGGGCAGTGACAATAGAGATCGAATTCTGTGC 2858
Nlrp1b_mRNA_KK CAGCACAAGACTCCACTTTGTAGTGACAAGGGCAGTGACAATAGAGATCGAATTCTGTGC  505

---                                                        N1b-R
Nlrp1b_mRNA_Ba CTTAGAAAAGTCAGGTGGGGTCTCTTTGGGATCCTGACATTGATAGGAGCCTGGCATCTC 3878
Nlrp1b_mRNA_KK CTTAGAAAAGTCAGGTGGGGTCTCTTTGGGATCCTGACATTGATAGGAGCCTGGCATCTC 1525

Nlrp1b_mRNA_Ba TTTTCCTGAGAGGCTTCATCCAGCGCTGGATGGAAACAGATACAGAGACCCACAGACAAC 3938
Nlrp1b_mRNA_KK TTTTCCTGAGAGGCTTCATCCAGCGCTGGATGGAAACAGATACAGAGACCCACAGACAAC 1585
```

**B**

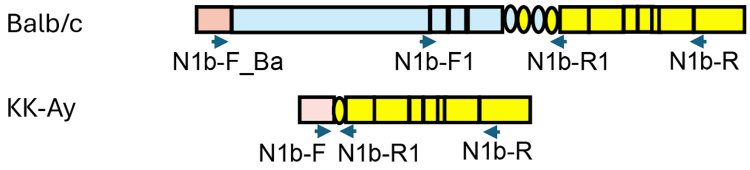


**C**

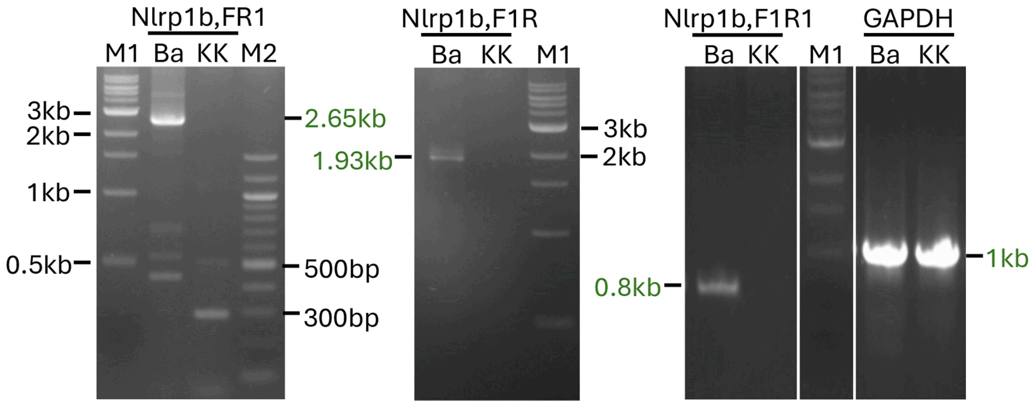

**Figure S5.** The full length *Nlrp1b* transcript is expressed in the bone marrow of Balb/c (Ba) mice, but not in KK-Ay (KK) mice. (**A**) Alignment of the *Nlrp1b* cDNA sequences of Balb/c and KK. The cDNA sequences were aligned to identify regions for primer design. The regions shown here were used for primer design. Ideally the same RT-PCR primers should be used both (Balb/c and KK-Ay) tissue, which is the case for the reverse primers N1b-R1 and N1b-R (highlighted green, the actual primer sequences are complimentary to the sequences shown). However, since the two exon 1 sequences are quite different (shown in red, with bolded fonts indicating the un-aligned areas), two different forward primers are used to generate amplicons starting from exon1 (highlighted yellow for N1b-F_Ba used for Balb/c samples and cyan for N1b-F used for KK-Ay samples). Primer N1b-F1 (highlighted yellow) targeting exons 2-3 is included to show that those exons are deleted in KK so no amplicons should be generated. The primer names are shown above or below the highlighted sequences (purple fonts). (**B**) A schematic illustration of the cDNA structure of the Balb/c and KK *Nlrp1b* transcripts. Exons are either rectangular or oval. The oval ones shown are exons 6 and 7, which are duplicated in Balb/c. Exons in yellow are identical or highly conserved between the two strains. The exons in blue are deleted in KK due to the 29 kb deletion (exons 2 to 6). The red (exon 1s) are not conserved. Exon numbering concurs with that of C57BL/6 *Nlrp1b*, which has 14 exons. (**C**) RT-PCR was performed on total RNA prepared from the bone marrow of KK-Ay and Balb/c mice. Three pairs of primers were used to amplify the *Nlrp1b* cDNA. The expected sizes of the Balb/c *Nlrp1b* amplicons are 2.65 kb (using N1b-F_Ba and N1b-R1, “Nlrp1b, FR1”), 1.93 kb (using primers N1b-F1 and N1b-R, “Nlrp1b, F1R”) and 0.8 kb (using primers N1b-F1 and N1b-R1, “Nlrp1b, F1R1”). The expected size of the *Nlrp1b* KK-Ay amplicon is 286 bp (if it expressed using N1b-F and N1b-R1 primers, “Nlrp1b, FR1”). No amplicons should be amplified when N1b-F1 was the forward primer. The RT-PCR results confirmed the effect of the KK deletion CNV . The band sizes are indicated in green and DNA ladder sizes are indicated in black. RT-PCR of GAPDH amplicons are shown as the control. The nature of the Balb/c (1.93kb and 0.8 kb) and KK (the 286 bp) amplicons and the *Gapdh* amplicons were confirmed by Sanger sequencing. M1, 1 kb DNA ladder; M2, 100 bp DNA ladder (New England Biolabs).

**Table S1. Characteristics of long-read sequencing (LRS) data obtained from 40 inbred strains.** For each strain, the table reports the Jackson Laboratory number, strain name, abbreviation used in the figures and main text, total HiFi sequencing output in gigabases (GB), sequencing coverage (×), total number of reads (in millions), mean read length, N50 read length, and median HiFi read quality value (QV). The N50 read length is included as a measure of read-length quality, defined as the length of the shortest read such that the longest reads collectively account for more than 50% of the total bases sequenced. Sequencing coverage was calculated by dividing the total HiFi yield by the estimated mouse genome size (2.73 Gb). Sequencing was performed on the PacBio Revio platform using the HiFi system.

| JAX Strain # | Strain | Abbrev | GB | Coverage (×) | # Reads (M) | Mean Read Length | N50 Read length (bp) | Median HiFi QV |
|---|---|---|---|---|---|---|---|---|
| #000463 | B10.D2-Hc1 H2d H2-T18c/nSnJ | B10D2 | 98.45 | 36.06 | 5.89 | 16722 | 18087 | Q31 |
| #000486 | MRL/MpJ | MRL | 96 | 35.16 | 6.17 | 15566 | 16825 | Q32 |
| #000550 | MOLF/EiJ | MOLF | 84.45 | 30.93 | 5.5 | 15345 | 16945 | Q33 |
| #000644 | SEA/GnJ | SEA | 90.93 | 33.31 | 7.14 | 12737 | 14857 | Q34 |
| #000646 | A/J | AJ | 85.12 | 31.18 | 4.69 | 18132 | 20255 | Q31 |
| #000648 | AKR/J | AKR | 94.11 | 34.47 | 5.63 | 16725 | 18181 | Q30 |
| #000651 | BALB/cJ | BALB | 108.88 | 39.88 | 6.67 | 16333 | 18050 | Q33 |
| #000653 | BUB/BnJ | BUB | 81.32 | 29.79 | 6.38 | 12741 | 14612 | Q34 |
| #000656 | CBA/J | CBA | 88.19 | 32.3 | 5.12 | 17230 | 19088 | Q31 |
| #000657 | CE/J | CE | 74.83 | 27.41 | 6.1 | 12266 | 13752 | Q32 |
| #000659 | C3H/HeJ | C3H | 87.2 | 31.94 | 5.53 | 15754 | 17701 | Q32 |
| #000664 | C57BL/6J | B6J | 87.65 | 32.11 | 5.05 | 17359 | 18965 | Q31 |
| #000665 | C57BL/10J | B10J | 92.54 | 33.9 | 5.76 | 16071 | 17362 | Q32 |
| #000668 | C57L/J | C57L | 92.2 | 33.77 | 5.92 | 15583 | 16863 | Q32 |
| #000669 | C58/J | C58J | 79.26 | 29.03 | 7.21 | 10997 | 13356 | Q35 |
| #000670 | DBA/1J | DBA1J | 109.58 | 40.14 | 6.58 | 16644 | 17989 | Q32 |
| #000671 | DBA/2J | DBA2J | 106.62 | 39.05 | 6.03 | 17694 | 19069 | Q32 |
| #000674 | I/LnJ | ILNJ2 | 91.93 | 33.67 | 6.65 | 13829 | 15595 | Q34 |
| #000676 | LP/J | LP | 89.05 | 32.62 | 5.49 | 16210 | 17873 | Q31 |
| #000677 | MA/MyJ | MAMy | 87.84 | 32.18 | 6.22 | 14119 | 15684 | Q33 |
| #000679 | P/J | PJ | 92.88 | 34.02 | 5.8 | 16150 | 17645 | Q34 |
| #000680 | PL/J | PL | 70.32 | 25.76 | 5.96 | 11803 | 13551 | Q33 |
| #000682 | RF/J | RF | 92.8 | 33.99 | 5.49 | 16893 | 18797 | Q32 |
| #000684 | NZB/BlNJ | NZB | 87.23 | 31.95 | 5.27 | 16550 | 18197 | Q30 |
| #000686 | SJL/J | SJL | 101.17 | 37.06 | 6.45 | 15679 | 17057 | Q33 |
| #000687 | SM/J | SMJ | 46.31 | 16.96 | 4.7 | 9864 | 11960 | Q35 |
| #000689 | SWR/J | SWR | 85.84 | 31.44 | 7.04 | 12194 | 14418 | Q35 |
| #000928 | CAST/EiJ | CAST | 79.22 | 29.02 | 5.94 | 13344 | 15657 | Q34 |
| #001058 | NZW/LacJ | NZW | 90.83 | 33.27 | 6.1 | 14884 | 15914 | Q31 |
| #001145 | WSB/EiJ | WSB | 97.26 | 35.63 | 6.03 | 16135 | 17391 | Q31 |
| #001146 | SPRET/EiJ | SPRET | 82.31 | 30.15 | 4.86 | 16936 | 18858 | Q33 |
| #001591 | RHJ/LeJ | RH2 | 81.86 | 29.99 | 7.67 | 10668 | 12523 | Q35 |
| #001800 | FVB/NJ | FVB | 97.39 | 35.67 | 6.07 | 16039 | 17481 | Q33 |
| #001976 | NOD/ShiLtJ | NOD | 87.21 | 31.95 | 4.92 | 17720 | 19586 | Q32 |
| #002050 | NOR/LtJ | NOR | 93 | 34.07 | 5.77 | 16114 | 17722 | Q32 |
| #002105 | NZO/HILtJ | NZO | 88.2 | 32.31 | 5.03 | 17538 | 19153 | Q31 |
| #002282 | BTBR T+ Itpr3tf/J | BTBR | 82.31 | 30.15 | 4.72 | 17433 | 18906 | Q32 |
| #002448 | 129S1/SvImJ | 129S1 | 84.49 | 30.95 | 5.16 | 16387 | 17795 | Q30 |
| #002468 | KK.Cg-Ay/J | KK | 88.64 | 32.47 | 5.4 | 16426 | 17674 | Q34 |
| #005314 | TALLYHO/JngJ | TH | 85.01 | 31.14 | 5.59 | 15211 | 16694 | Q31 |
| **Total** | | | **3540.45** | | **233.69** | | | |
| **Mean** | | | **88.51** | **32.42** | **5.84** | **15301** | **16952** | |

**Table S2. The PCR primers used to validate 10 deletion CNVs.** The flanking primers used to amplify a fragment from strains with CNV deletion alleles are selected because the deletion results in the primers being located in proximity to each other. As a control, internal primers are used to amplify fragments from C57BL/6J tissue, which does not have the deletion allele.

| CNV | ID_T2T | Strains with Deletion | Primers Flanking the CNV Deletion | | | Internal Primers for B6J | | |
|---|---|---|---|---|---|---|---|---|
| | | | Primers | Primer Sequence | Expected PCR Fragment | Internal Primers | Internal Rev Primer Sequence | Expected PCR Fragment |
| CNV1 | chr3-31102768-DEL-13674 | 129S1,BTBR,C57L,DBA2J,MAMy | CNV1-F | TTCTATGGTGTACTTGACGTCCCT | 725 bp | CNV1-R1 | GTAGATGGAAGGAGTGGACTTGGT | 577 bp |
| | | | CNV1-R | GCTGTCTCCTAAATAGAATCTGGGTCTG | | CNV1-F1 | AACATTTCTAATATTCTGCATCCCTGGA | 901 bp |
| CNV2 | chr14-108331180-DEL-66424 | NZB,B10J | CNV2-F | TGTAAGAACATGCCAGAGCTATTGCT | 353 bp | CNV2-R1 | CCACTGCTTGCTTGACAAAATACCA | 602 bp |
| | | | CNV2-R | TGTTAATTTACTGTATCAGGAGCCAGGA | | CNV2-F1 | AGAGCCTCAGCACTATTCTTCTTTTGA | 549 bp |
| CNV3 | chr5-50693484-DEL-134404 | NOD,NOR | CNV3-F | ATGAGTTGTCAGGATCAATGGTGGA | 949 bp | CNV3-R1 | GCACAAAAAGAGTTCTGGTTGGCT | 647 bp |
| | | | CNV3-R | TCCAAGCAGAATGAATACACCAGGT | | CNV3-F1 | AGAATCTTCTAATGTGATACTCACGTCTCT | 701 bp |
| CNV4 | chr5-61410898-DEL-11558 | KK | CNV4-F | TAGCATATCTAGGTCCCTTTGAGACA | 1113 bp | CNV4-R1 | AGGTGAGCGCAACAAATAATTATCAA | 904 bp |
| | | | CNV4-R | TCCTTCCATCTCAGTTCCAGAATATGT | | CNV4-F1 | AAATGGGTAAACTCTCTAGATGGTTGTT | 401 bp |
| CNV5 | chr6-81578193-DEL-34289 | AJ,BTBR | CNV5-F | TCCAGTTGAAAGTCACAGGTATCTTTAC | 735 bp | CNV5-R1 | TGTTTGTAGGCCATCTGACTTCTGT | 919 bp |
| | | | CNV5-R | GTATGTGTTACAGGGTGTTCCAGATCA | | CNV5-F1 | AGAGTTTATTACTGCAGCAGCGCT | 254 bp |
| CNV6 | chr7-125131880-DEL-18679 | SJL | CNV6-F | AGGTGCCGTTAATCCAGGTCAAGAT | 344 bp | CNV6-R1 | TGTTTTACTCAAAGCCACCTGAAGTT | 828 bp |
| | | | CNV6-R | TCAGGCAGAAAGAAGGACCTAGCAA | | CNV6-F1 | GTGAAGATTACCCACTTCCATGTTCA | 625 bp |
| CNV7 | chr7-155981379-DEL-36228 | C58J,LP,MAMy,PJ,SMJ | CNV7-F | CCAAGCTGCAGAAGAAGCCAAAC | 1118 bp | CNV7-R1 | ACGATTCCAACAATAGTTCATGCCTCT | 383 bp |
| | | | CNV7-R | CAAGGAATTGGCCCCTCCCA | | CNV7-F1 | GAGACGTGGCTGAAGGTAACCAT | 1119 bp |
| CNV8 | chr8-68531275-DEL-15963 | C58J,LP,PJ,SEA | CNV8-F | TGCAGAGAGGTGTAACAGGAAGCT | 686 bp | CNV8-R1 | TGGACCTAGACCTCTATGAATATATGTCA | 1000 bp |
| | | | CNV8-R | GTAATGGCTTTCCTTATAGCTTTGGCA | | CNV8-F1 | CATTTGAATGCTGATGATATATCAATCCCT | 593 bp |
| CNV9 | chr14-94842485-DEL-10521 | AJ,BALB,BUB,C3H,CBA,CE,FVB,SEA,SJL,SMJ,SWR,TH | CNV9-F | GTTCACAGCTCTAAGTCAGGACAGT | 575 bp | CNV9-R1 | GGGGCTCGACGTTTTTAAATTTCT | 498 bp |
| | | | CNV9-R | AGTCTATGAACATTTGTACTTCCAAGTCT | | CNV9-F1 | TGACTCTGTCACTGAAACTCGATCA | 508 bp |
| CNV10 | chr15-70022454-DEL-15124 | AKR,BTBR,ILNJ2,NZO | CNV10-F | ATTTATCCCTTCATAGGTCTCTGGTTCA | 704 bp | CNV10-R1 | GTGCATTCTTTCTGGCGAGATGGT | 368 bp |
| | | | CNV10-R | CAGTCAGTTTATACTATTACAGAGCACCT | | CNV10-F1 | AAGTACAGTTGGAGACGTGTACCA | 562 bp |

**Table S3. Association of human *NLRP1* CNV, INDEL, and SNP alleles with quantitative traits associated with the metabolic syndrome.** The association statistics were obtained from the UK Biobank–AstraZeneca PheWAS Portal and FinnGen release 13 PheWeb databases. All variants are located entirely within the human *NLRP1* gene body (GRCh38, chr17:5,499,415–5,619,424). The variant class (CNV, INDEL, or SNP) and identifier (Variant/rsID) and database are indicated; and NA indicates that no rsID was available. CNV sizes were calculated from the reported GRCh38 breakpoints. Annotations, phenotype names, effect estimates, *P* values, and sample sizes are those reported by the respective database. The FinnGen Body Mass Index (BMI) and weight traits were inverse-rank normalized.

| Variant Class | Variant / rsID | Position | Annotation | Dataset | Phenotype | Effect | *P* value | Sample Size |
|---|---|---|---|---|---|---|---|---|
| CNV | L:17-5518093-5539753 | chr17:5,518,093–5,539,753; 21.66 kb | Deletion | UKB–AstraZeneca | Glycated hemoglobin, HbA1c | 1.044 | $1.67\times10^{-3}$ | 422,544 |
| CNV | L:17-5518093-5539753 | chr17:5,518,093–5,539,753; 21.66 kb | Deletion | UKB–AstraZeneca | HDL cholesterol | -1.077 | $3.42\times10^{-3}$ | 386,995 |
| CNV | L:17-5518093-5539753 | chr17:5,518,093–5,539,753; 21.66 kb | Deletion | UKB–AstraZeneca | Apolipoprotein A | -1.205 | $1.22\times10^{-3}$ | 384,811 |
| INDEL | rs200485068 | chr17:5,511,049 CA>C | 1-bp deletion | FinnGen R13 | BMI, inverse-rank normalized | 0.029 | $8.41\times10^{-9}$ | 362,216 |
| INDEL | rs200485068 | chr17:5,511,049 CA>C | 1-bp deletion | FinnGen R13 | Weight, inverse-rank normalized | 0.030 | $3.68\times10^{-10}$ | 369,975 |
| INDEL | NA | chr17:5,553,855 C>CT | 1-bp insertion | FinnGen R13 | BMI, inverse-rank normalized | 0.027 | $4.98\times10^{-8}$ | 362,216 |
| INDEL | NA | chr17:5,553,855 C>CT | 1-bp insertion | FinnGen R13 | Weight, inverse-rank normalized | 0.028 | $2.10\times10^{-9}$ | 369,975 |
| SNP | rs2301582 | chr17:5,532,943 C>T | Missense | UKB–AstraZeneca | Insulin-like growth factor 1 (IGF-1) | -0.015 | $3.09\times10^{-12}$ | 432,720 |
| SNP | rs2301582 | chr17:5,532,943 C>T | Missense | UKB–AstraZeneca | Insulin-like growth factor 1 (IGF-1) | -0.025 | $5.69\times10^{-10}$ | 432,720 |
| SNP | rs2301582 | chr17:5,532,943 C>T | Missense | FinnGen R13 | BMI, inverse-rank normalized | -0.009 | $1.98\times10^{-5}$ | 362,216 |
| SNP | rs2301582 | chr17:5,532,943 C>T | Missense | FinnGen R13 | Weight, inverse-rank normalized | -0.009 | $1.71\times10^{-5}$ | 369,975 |
| SNP | rs11651270 | chr17:5,521,757 T>C | Missense | UKB–AstraZeneca | Insulin-like growth factor 1 (IGF-1) | -0.013 | $1.14\times10^{-9}$ | 432,759 |
| SNP | rs11651270 | chr17:5,521,757 T>C | Missense | UKB–AstraZeneca | Insulin-like growth factor 1 (IGF-1) | -0.020 | $1.33\times10^{-8}$ | 432,759 |
| SNP | rs11651270 | chr17:5,521,757 T>C | Missense | FinnGen R13 | BMI, inverse-rank normalized | -0.009 | $3.24\times10^{-5}$ | 362,216 |
| SNP | rs11651270 | chr17:5,521,757 T>C | Missense | FinnGen R13 | Weight, inverse-rank normalized | -0.008 | $3.71\times10^{-5}$ | 369,975 |
